# Ratiometric Fluorescent Protein Biosensors Reveal Citrate Dynamics and Cellular Heterogeneity

**DOI:** 10.64898/2026.04.16.718871

**Authors:** Saaya Hario, Norito Tamura, Bibi Safeenaz Alladin-Mustan, Syed Musa Ali, Matthew S. Macauley, Yi Shen, Robert E. Campbell, Ina Huppertz, Kei Takahashi-Yamashiro

**Author notes:** Corresponding authors: Ina Huppertz, Max Planck Institute for Biology of Ageing, Joseph-Stelzmann-Str. 9b, 50931 Cologne, Germany, Kei Takahashi-Yamashiro, Department of Chemistry, Graduate School of Science, The University of Tokyo, 7-3-1 Hongo, Bunkyo-ku, Tokyo 113-0033, Japan. These authors contributed equally to this work (Saaya Hario, Norito Tamura).

## Abstract

Citrate is a central intermediate metabolite linking the tricarboxylic acid cycle and lipid biosynthesis. Tools for monitoring of spatiotemporal citrate dynamics are critical for getting a better understanding of cellular metabolism. Here, we develope genetically encoded excitation ratiometric biosensors for citrate, based on our previous intensiometric green fluorescence protein-based citrate biosensor, Citron1. We find that a single mutation in the Citron1 chromophore-forming tripeptide provided an excitation ratiometric response. Further rounds of directed evolution yield highly responsive variants, exhibiting citrate-dependent fluorescence changes between two excitation peaks. When expressed in mammalian cells, these biosensors enable citrate dynamics to be monitored in both the cytosol and mitochondria. Comparative analysis across multiple human breast cancer cell lines uncovers cell line-specific differences in citrate levels and their heterogeneity, which could be linked to their malignancy. Furthermore, flow cytometry-based measurements in mouse embryonic stem cells demonstrate the proteomics signatures underlying the population-level variability in citrate concentrations and citrate rewiring during stem cell differentiation. Together, these results show that these excitation ratiometric citrate biosensors enable quantitative, compartment-resolved, and population-scale analysis of cellular metabolism.

## Introduction

Metabolism is highly controlled by a myriad of metabolite interconnections and plays a critical role in maintaining cellular function. Metabolic rewiring has been recognized as a critical event during development, in sustaining cellular homeostasis, and in the progression of diseases including cancer^1^. Recent advances in mass spectrometry (MS)-based metabolomics and lipidomics techniques have made it possible to detect the levels of numerous metabolites in biological contexts and have begun to reveal the complexity of metabolism in mammalian tissue^2^. However, the spatiotemporal dynamics of cellular metabolites remains largely unknown, and it is challenging to evaluate metabolite levels at the single-cell level.

Citrate is an important intermediate substrate of the tricarboxylic acid (TCA) cycle in mitochondria^3^. It is produced from acetyl-CoA and oxaloacetate by citrate synthase (CS), then converted into isocitrate via cis-aconitate by aconitase 2 (ACO2). In the cytosol, citrate is converted to acetyl-CoA by ATP citrate lyase (ACLY). Acetyl-CoA further acts as a building block for fatty acid and sterol biosynthesis. In neuronal cells, citrate is a metabolic precursor for the biosynthesis of glutamate, γ-aminobutryate (GABA) and acetylcholine^4^. Citrate is transported between the cytosol and the mitochondria through the mitochondrial citrate carrier, SLC25A1, also known as citrate/isocitrate carrier (CIC)^3,5^. While CIC exports citrate from mitochondria to the cytoplasm, it simultaneously imports malate from the cytoplasm to the mitochondrial matrix. CIC is essential for the exit of embryonic stem cells (ESCs) from pluripotency and supports embryonic development; loss of CIC causes proliferation defects and senescence^6^. In mouse ESCs, CIC enables a metabolic shift to a non-canonical TCA cycle when cells transition from the naïve to a primed state^7^. Furthermore, the expression levels of CIC are linked to inflammation and increased by the stimulation of proinflammatory cytokines, such as tumor necrosis factor (TNF)-α and interferon (IFN)-γ^8^. It has also been reported that CIC expression is upregulated in cancer cells and the inhibition of CIC could suppress proliferation of cancer cell lines^9^. These observations suggest that citrate concentration levels are closely related to disease progression.

Another important citrate transporter is the Na^+^-dependent citrate transporter (NaCT), also known as SLC13A5 or plasma membrane citrate transporter (pmCiC), which is responsible for the import and export of citrate across the plasma membrane^4^. NaCT is expressed specifically in the liver, testes and brain^4,10^. Mutations in SLC13A5 cause citrate transport dysfunction and can lead to epilepsy^11,12^. In cancer cells, the cellular citrate concentration is increased to enable rapid proliferation, and the uptake of citrate through NaCT contributes to metastasis^13,14^. Conversely, reduced citrate levels were also observed in cancer cells and exogenous citrate treatment inhibited their cell proliferation^15^. Understanding citrate dynamics will help us to better assess metabolic homeostasis and interpret diseases.

Genetically encoded biosensors based on fluorescent proteins (FPs) are powerful tools for a real-time visualization of ions and molecules inside cells and tissues^16^. Recently, several high performance FP-based biosensors have been developed and applied for the monitoring of metabolites such as citrate, lactate and pyruvate^17–19^. Single FP-based biosensors can be categorized into two types: intensiometric and ratiometric. In both types of biosensors, the underlying mechanism relies on a ligand binding-induced conformational change in the sensor domain (SD). These conformational changes affect the chromophore environment leading to a change in fluorescence properties. Many single FP-based intensiometric biosensors exhibit a ligand-dependent change in fluorescence intensity, but fluorescence intensity is also influenced by biosensor expression levels and imaging conditions, which can limit quantitative interpretation and long-term monitoring. On the other hand, ratiometric biosensors are generally more suitable for quantitative analysis because the ratio of two peaks is, ideally, independent of biosensor concentration and less sensitive to small fluctuations (e.g., in focus, light intensity, background, etc.), that affect both peaks equally.

Classically, an inherent ratiometric response has been seen as a major advantage of Förster resonance energy transfer (FRET)-based biosensors, which are composed of two FPs and a SD. Another type of ratiometric biosensors, excitation-ratiometric biosensors, consist of a single FP and a SD and is somewhat more suitable for multicolor imaging^20^ because its excitation and emission bands span a smaller range of wavelengths. Notably, by fusing a second FP, single FP-based intensiometric biosensors can be converted into ratiometric biosensors, but at the cost of expanding the range of wavelengths required for a single biosensor.

Our group has previously developed high performance single green FP (GFP)-based intensiometric biosensors for citrate, named Citron1 and Citroff1^19^. However, a ratiometric version is not available yet. In this study, we developed high performance GFP-based excitation ratiometric citrate biosensors, which we describe as the ratioCitron1 series, based on the original Citron1 biosensor^19^. We evaluated the ratioCitron1 biosensor series in various types of mammalian cells, using not only fluorescence microscopy for single cell measurements, but also flow cytometry to provide population-based measurements. As a result, we revealed the cancer cell line specific citrate levels and their heterogeneity, which could be related to their malignancy. The combination of ratioCitron1 biosensors and omics analysis demonstrated the proteomics signatures linked to the citrate levels in mESCs. We also analyzed the involvement of citrate levels in the differentiation of mESCs. This study paves the way to their wide range of biological applications.

## Results

### Development of an excitation-ratiometric biosensor for citrate

In our previous work on developing intensiometric biosensors for citrate, the citrate-binding periplasmic domain (CitAP) of *Klebsiella pneumoniae* sensor histidine kinase (SHK) CitA was used as a SD^19^. CitAP was inserted into GFP, and linker optimization was used to produce prototypes with direct- and inverse-fluorescence responses upon citrate binding. After directed evolution, which is an iterative process of random mutagenesis and screening, Citron1 and Citroff1 were developed^19^. An initial Citron1-derived prototype with an excitation ratiometric response was discovered while exploring the effect of introducing mutations in the chromophore. Specifically, we found that introduction of a T65A mutation in the chromophore-forming tripeptide produced an excitation-ratiometric response (**Supplementary Fig. 1**). To improve this response, characterized by the parameter Δ*R*/*R* (where Δ*R*/*R =* (*R*_on_ – *R*_off_)/*R*_off_, *R* = (*F* with excitation ∼470 nm and emission ∼510 nm)/(*F* with excitation ∼400 nm and emission ∼510 nm), “on” = citrate binding, and “off” = citrate non-binding), 11 rounds of directed evolution were conducted (**Fig. 1A**) with library creation being done by error-prone PCR of the whole coding gene. The best variant identified in the final round had 16 mutations, with 14 in the GFP region and 2 in the CitAP region, in addition to the original T65A mutation (**Fig. 1B**, **Supplementary Fig. 2**). *In vitro* characterization of this final variant, which was temporarily designated as ratioCitron1, revealed that it has a primary excitation peak at 398 nm in the absence of citrate and at 492 nm in the presence of citrate, and excitation at either peak leads to an emission peak at ∼513 nm (**Fig. 1C**, **Table 1**). The emission peak excited by a wavelength of 400 nm shows a higher value in the absence of citrate than its presence (**Fig. 1C**). Conversely, the emission peak excited by a wavelength of 450 nm shows a lower peak in the absence of citrate than in its presence (**Fig. 1C**). Thus, the ratio of these two peaks is dependent on the citrate concentration. The Δ*R*/*R* value of the final variant was 25.6 upon the addition of 90 mM citrate, when measured using purified protein. The absorbance spectra revealed peaks at around 391 nm and 497 nm, corresponding to the protonated and anionic states of the chromophore, respectively (**Fig. 1D**).

**Figure 1.**
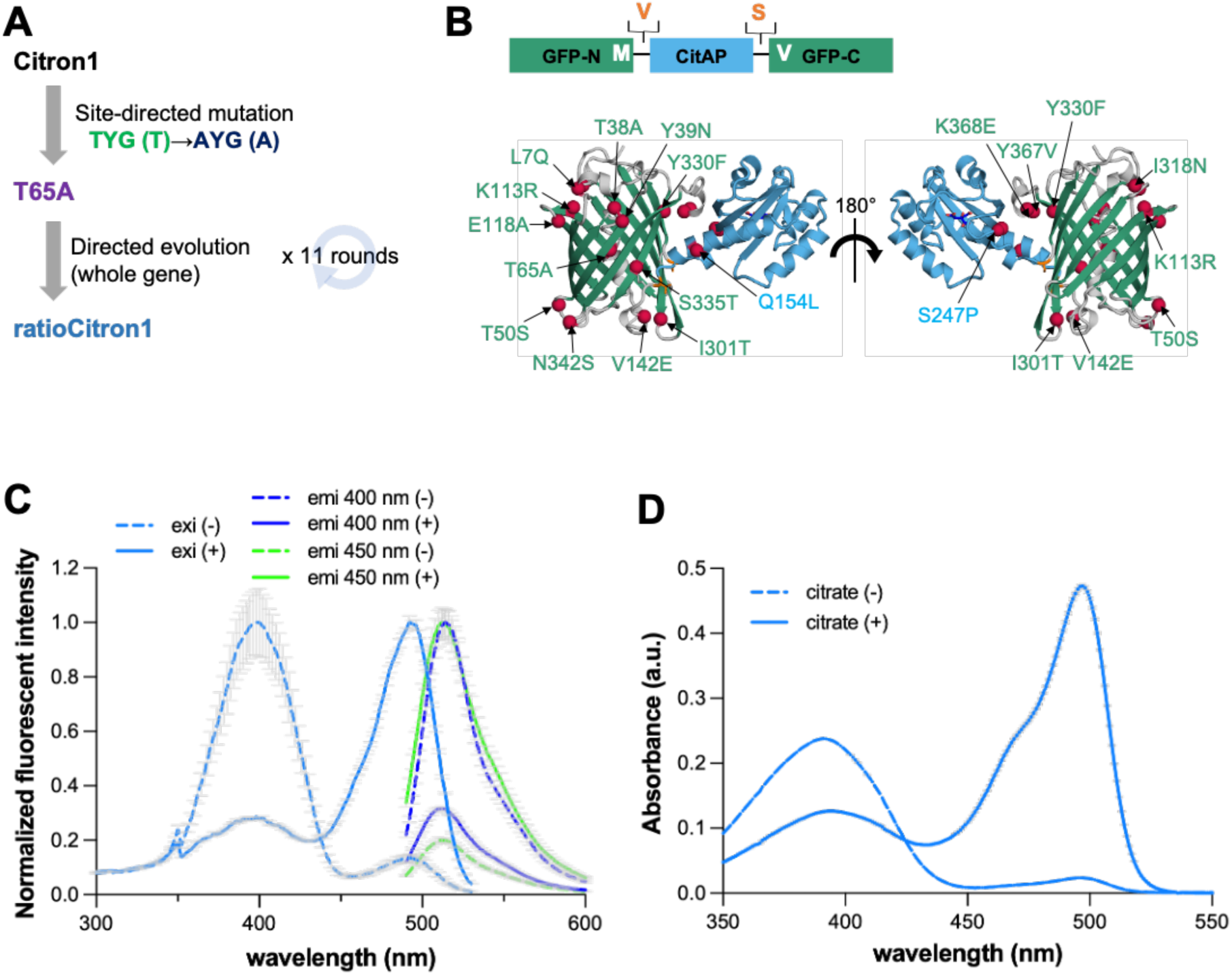
Development of ratioCitron1. **A.** Development scheme of ratioCitron1. **B.** A schematic representation (upper) and an AlphaFold model^21^ (lower) of ratioCitron1. GFP is shown in green, CitA is shown in cyan, and introduced mutations are in red. **C.** Excitation (light blue lines) and emission spectra (blue and green lines) of ratioCitron1 in the absence (dotted lines) and presence (solid lines) of citrate (90 mM). *n* = 3 technical replicates, mean ± SD. **D.** Absorbance spectra of ratioCitron1 in the absence (dotted line) and presence (solid line) of citrate (90 mM). *n* = 3 technical replicates, mean ± SD.

**Table 1.** Photophysical properties of ratioCitron1 variants.

| variants | $\Delta R/R$ | Apparent $K_d$ [μM] | $pK_a$ | ligand (-/+) | excitation peak (nm) | emission peak (nm) | |
| --- | --- | --- | --- | --- | --- | --- | --- |
|  |  |  |  |  |  | exc 400 nm | exc 450 nm |
| ratioCitron1Low | 25.6 | 7300 | 10 | (-) | 398 | 514 | - |
|  |  |  | 7.8 | (+) | 492 | - | 512 |
| ratioCitron1High | 26.0 | 420 | 10 | (-) | 398 | 514 | - |
|  |  |  | 6.8 | (+) | 494 | - | 512 |
| control_ratioCitron1Low | -0.08 | - | - | (-) | 490 | 510 |  |
|  |  |  | - | (+) | 490 |  |  |
| control_ratioCitron1High | -0.10 | - | - | (-) | 490 | 512 |  |
|  |  |  |  | (+) |  |  |  |

### Development of high affinity and control variants

Physiological citrate concentrations range from hundreds of μM to several mM^4,22^. When ratioCitron1 was expressed in HeLa cells, and the cells were treated with digitonin to permeabilize the membrane, no substantial response was observed for 1 mM of exogenously added citrate, but a substantial response was observed with 10 mM citrate (**Supplementary Fig 3A**). This result suggested that ratioCitron1 had an apparent *K*_d_ in the low mM range. In an effort to develop a higher affinity variant, we conducted site-directed and site-saturation mutagenesis on the amino acid residues reported to contribute to the binding to citrate^19^ (**Fig. 2A**, labeled in blue), and screened the resulting variants in terms of response and apparent *K*_d_ for citrate. This effort did not lead to any promising variants with higher affinity. Next, we aligned the amino acid sequences of CitA in ratioCitron1, Citron1, Citroff1, and original CitAP (**Supplementary Fig. 2**). Since ratioCitron1 exhibited lower affinity compared to Citron/off1 and original CitAP, we hypothesized that there could be the mutations introduced during the development decreasing the affinity. Therefore, we reversed selected mutations that had been introduced into the CitA domain during the development of Citron1 and in this work (**Fig. 2A**, labeled in magenta and red). The G236S (**Fig. 2A**, labeled in red) variant, when expressed in HeLa cells, showed higher affinity while maintaining a Δ*R*/*R* value indicative of a response to the 1 mM exogenously added citrate (**Supplementary Fig. 3B**). Thus, this residue was further randomized and the resulting variants were screened to identify higher affinity variants. This effort led to the discovery of the G236H variant with an apparent *K*_d_ of less than 1 mM. The G236H variant was designated as ratioCitron1High (**Supplementary Fig. 3C‒D**), and the variant previously referred to as ratioCitron1 was designated as ratioCitron1Low.

**Figure 2.**
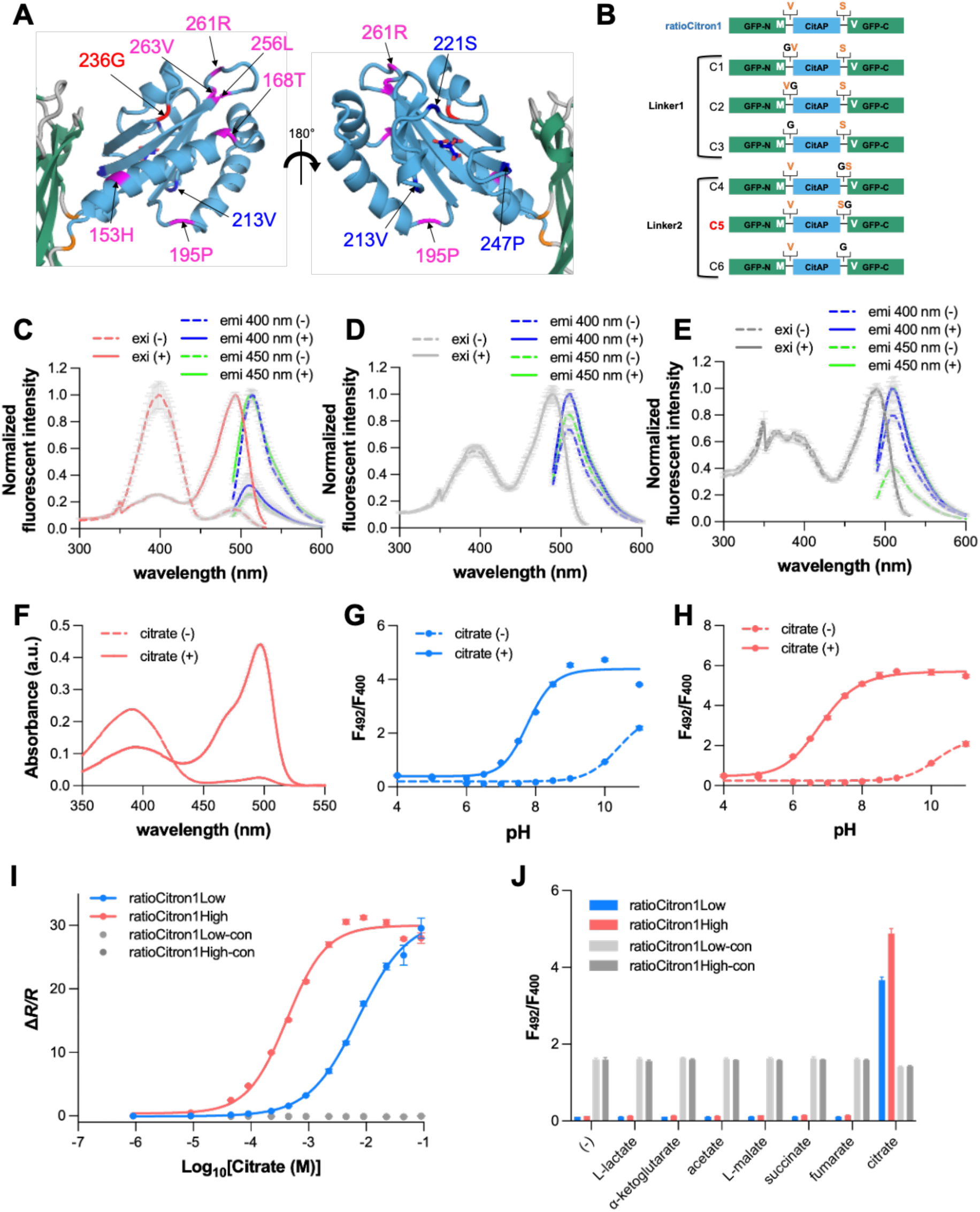
Development and *in vitro* characterization of high affinity and control variants. **A.** Mutations tested to develop a high affinity variant of ratioCitron1. The key residues for citrate binding are shown in blue. The residues in CitA where mutations are inserted in the development of Citron1 and ratioCitron1 are shown in magenta and 236G is shown in red. **B.** Design of control variants of ratioCitron1 variants. The linker residues (V and S) are in orange and the glycine insertion patterns (C1-6) are shown. **C-E.** Excitation and emission spectra of ratioCitron1High (**C**) and control variants (ratioCitron1Low-con (**D**) and ratioCitron1High-con (**E**)) in absence and presence of citrate. *n* = 3 technical replicates, mean ± SD. **F.** Absorbance spectra of ratioCitron1High in the absence and presence of citrate. **G-H.** pH titrations of ratioCitron1Low (**G**) and ratioCitron1High (**H**). **I.** Citrate titration of all ratioCitron1 variants. *n* = 3 technical replicates, mean ± SD. **J.** Ligand specificity of ratioCitron1 variants. *n* = 3 technical replicates, mean ± SD.

We next engineered control variants, which fluoresce but do not substantially change Δ*R*/*R* values upon the binding to citrate, for both ratioCitron1Low and ratioCitron1High. Using these control biosensors enables us to confirm the fluorescence change is caused by changes in citrate concentration, and not by other factors. To engineer the control variants while maintaining the same affinity for citrate, we constructed six prototypes by substituting residues for glycine, or adding an additional glycine, in one of the linkers in ratioCitron1Low (**Fig. 2B**). We hypothesized that these glycine mutations could abolish the key interactions between the SD and the chromophore, which would result in appropriate control variants. Potential mutants were screened in the context of both ratioCitron1Low and ratioCitron1High. Ultimately, we found that insertion of a glycine at the C-terminal end of the second linker (C5 in **Fig. 2B**) produced control variants for ratioCitron1Low and ratioCitron1High with the desired properties. We designated these control constructs as ‘ratioCitron1Low-con’ and ‘ratioCitron1High-con’, respectively.

### *In vitro* characterization of ratioCitron variants

The emission and excitation spectra of ratioCitron1High revealed that the excitation peak is at 398 nm in the absence of citrate and at 494 nm in the presence of citrate. Both excitations lead to an emission peak at around 513 nm (**Fig. 2C**, **Table1**). The control variants exhibit excitation peaks at 490 nm in both the absence and presence of citrate (**Fig. 2D‒E**). The absorbance spectra of ratioCitron1High reveals peaks at 391 nm in the absence of citrate, and at 497 nm in its presence (**Fig. 2F**). The p*K*_a_ values of ratioCitron1Low were 10 and 7.8 in the absence and presence of citrate, respectively, while ratioCitron1High showed p*K*_a_ values of 10 and 6.8 in the absence and presence of citrate (**Fig. 2G‒H**). The control variants showed similar absorbance spectra and pH titration curves in both the absence and presence of citrate (**Supplementary Fig. 4A‒D**). Notably, the citrate titration curves of the ratioCitron1 variants showed that these biosensors could be used to detect citrate concentrations ranging from the tens of μM to the hundreds of mM. In contrast, the control variants did not respond to concentrations of citrate as high as 100 mM. The *K*_d_ values were 7.3 mM and 420 μM for ratioCitron1Low and ratioCitron1High, respectively (**Fig. 2I**). The ratioCitron1 variants also showed good specificity, and the control variants did not respond to any other metabolites (**Fig. 2J**). The photophysical properties of all the ratioCitron1 variants are summarized in **Table1**.

### Imaging characterization of ratioCitron1 variants in mammalian cells

The ratioCitron1 series (i.e., ratioCitron1Low, ratioCitron1Low-con, ratioCitron1High, and ratioCitron1High-con), were expressed in HeLa cells and their responses to intracellular citrate concentration changes were determined. HeLa cells were permeabilized with digitonin and 20 mM citrate was added exogenously. **Fig. 3A‒B** shows pseudo-colored images of green fluorescent emission, with excitation at 405 nm and 470 nm, before and after the addition of 20 mM citrate in HeLa cells expressing ratioCitron1Low (**Fig. 3A**) and ratioCitron1High (**Fig. 3B**). Time courses of Δ*R*/*R* and Δ*F*/*F* of ratioCitron1Low (**Fig. 3C**) and ratioCitron1High (**Fig. 3D**) upon the addition of citrate indicate that both variants respond to changes in intracellular citrate concentration. **Fig. 3E** shows the responses of all ratioCitron1 variants to 0.1 mM, 1 mM, and 10 mM citrate in digitonin-treated HeLa cells. RatioCitron1High responded to 1 mM citrate and both ratioCitron1High and ratioCitron1Low responded to 10 mM citrate, while control variants showed negligible change in Δ*R*/*R* values.

**Figure 3.**
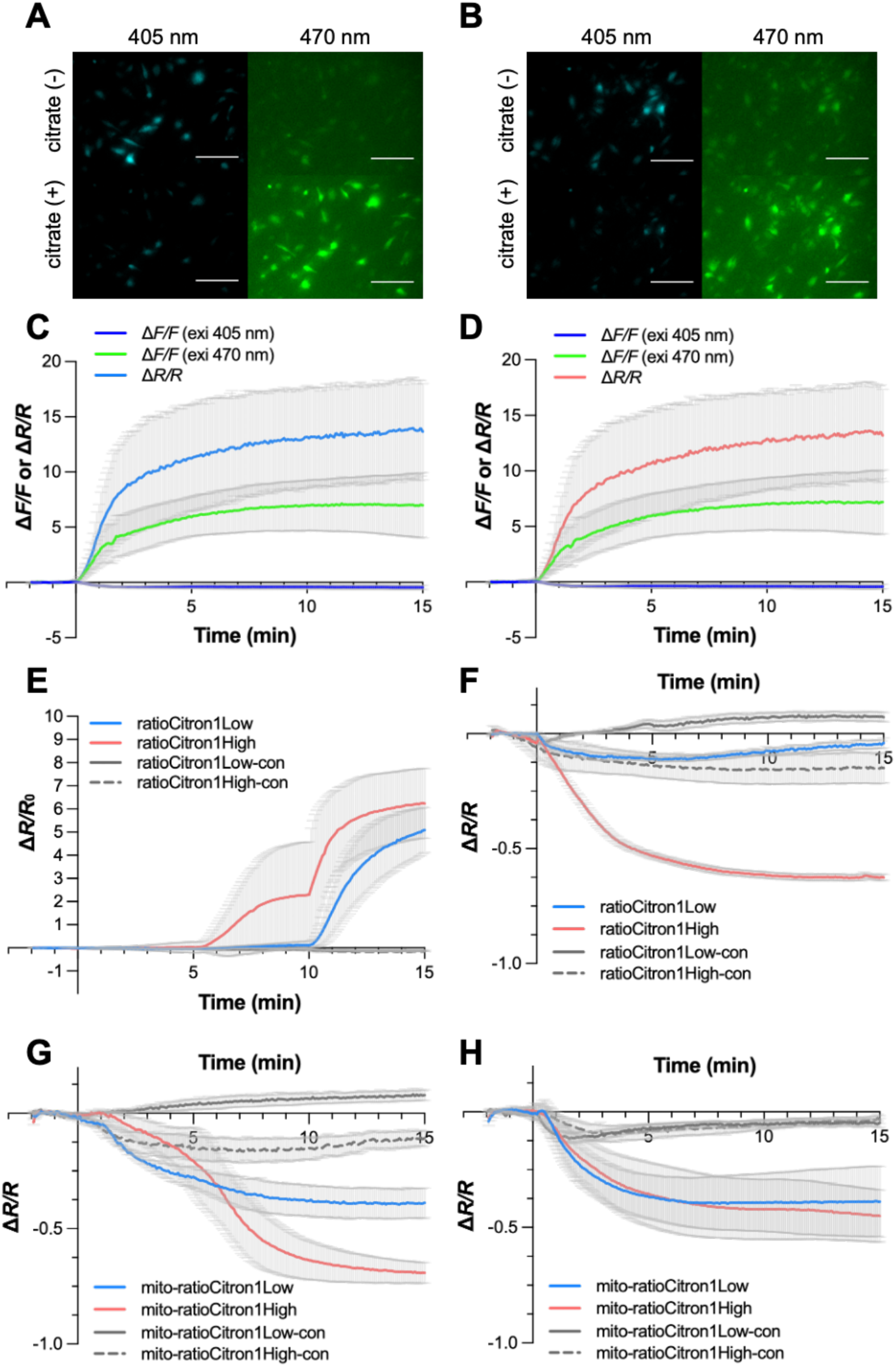
RatioCitron1 tracks citrate dynamics in HeLa cells. **A-B.** Pseudo-colored images of HeLa cells expressing ratioCitron1Low (**A**) and ratioCitron1High (**B**) in cytosol before and after addition of citrate. Scale bar: 100 µm. **C-D.** Δ*R*/*R* and Δ*F*/*F* values of ratioCitron1Low (**C**) and ratioCitron1High (**D**) versus time upon the addition of citrate. HeLa cells were incubated with 4 μM digitonin for 10 mins before the experiment and a final concentration of 20 mM citrate was added at t = 0 (ratioCitron1Low *n* = 16, ratioCitron1High *n* = 19, mean ± SD). **E.** Time courses of Δ*R*/*R*₀ of ratioCitron1 variants (ratioCitron1Low (*n* = 20), ratioCitron1High (*n* = 10), ratioCitron1Low-con (*n* = 5), ratioCitron1High-con (*n* = 5)) expressed in HeLa cells upon the citrate titration. HeLa cells expressing ratioCitron1 variants were treated with digitonin and citrate titration was carried out. The final concentration of citrate is 100 μM at 0 min, 1 mM at 5 min, 10 mM at 10 min (mean ± SD). **F-G.** Δ*R*/*R* values of ratioCitron1 variants expressed in cytosol (**F**) and in mitochondria (mito-ratioCiton1) (**G**) of HeLa cells upon the addition of the ACLY inhibitor of BMS-303141. **H.** Δ*R*/*R* values of ratioCitron1 variants expressed in mitochondria (mito-ratioCitron1) of HeLa cells upon the addition of MPC inhibitor UK-5099 (final concentration: 25 μM). Δ*F/F* = (*F*_on_ - *F*_off_)/*F*_off_, and *F* with excitation 405 nm or 470 nm and emission 518/45 nm. Δ*R*/*R* = (*R*_on_ - *R*_off_)/*R*_off_, and *R* = (*F* with excitation 470 nm and emission 518/45 nm) / (*F* with excitation 405 nm and emission 518/45 nm).

We tested several pharmacological applications using the ratioCitron1 variants in the context of HeLa cells. We first tested BMS 303141, which is an inhibitor of adenosine triphosphate (ATP)-citrate lyase (ACLY), an enzyme generating acetyl-CoA from citrate. Time-courses of Δ*R*/*R* of ratioCitron1 variants expressed in the cytosol (**Fig. 3F**) and in mitochondria (**Fig. 3G**) showed a decrease in intracellular citrate concentration, which is similar to what we observed with Citron1^19^.

We also tested UK-5099, which is an inhibitor of the mitochondrial pyruvate carrier (MPC), which blocks the transport of pyruvate from the cytosol to the mitochondria that fuels the TCA cycle. For this application, the ratioCitron1 variants were expressed in mitochondria. Both ratioCitron1Low and ratioCitron1High showed a decrease in Δ*R*/*R* upon the addition of UK-5099. This is consistent with a disrupted pyruvate import into the mitochondria, which leads to a reduction in citrate levels (**Fig. 3H**). These data demonstrated that the ratioCitron1 series can detect the citrate dynamics in mammalian cells.

### Monitoring citrate levels in various human breast cancer cell lines

By leveraging the advantages of the ratiometric biosensors, the citrate levels were compared between different cell lines. This is not suitable for single-FP based intensiometric biosensors. Several human breast cancer cell lines (MCF-7, BT549, T47D, ZR-75-1) in addition to HEK293T cells were transfected with the series of ratioCitron1 or non-responsive control biosensors. The cell images were captured using lasers with wavelengths of 385 nm and 475 nm. The emission ratio of each excitation wavelength (F_475_/F_385_) was higher in mitochondria in all cell lines compared to the cytosol (**Fig. 4A‒B**), meaning that the citrate concentration is higher in mitochondria. A low *K*_d_ variant, ratioCitron1High, showed a higher F_475_/F_385_ than the one from ratioCitron1Low, consistent with their *K*_d_ values. Interestingly, the F_475_/F_385_ values, i.e. the citrate levels, are heterogenous in some cell lines. For example, MCF-7 and T47D cells show high heterogeneity of citrate concentration in both cytosol and mitochondria. On the other hand, ZR-75-1 and BT549 cells show less heterogeneity and the citrate concentration is lower compared to the other 3 cell lines. Although both ratioCitron1 variants can detect the heterogeneity of citrate levels, ratioCitron1High is particularly suitable for the cytosolic citrate detection. This also suggests the importance of picking the right *K*_d_ variant for more precise analysis. Using control biosensors, we ruled out other factors as the source of this effect (**Supplementary Fig. 5**). These results demonstrated that the ratioCitron1 variants enable us to compare the citrate levels among different cancer cell lines. In addition, we found heterogeneous citrate levels due to the large sample size, highlighting the importance of single-cell metabolite analysis to understand correlations with biological events. This heterogeneity and variation between cancer cell lines would likely be missed by population-based approaches such as metabolomics. For future analysis, it will be interesting to see how the citrate concentration scales with cancer malignancy.

**Figure 4.**
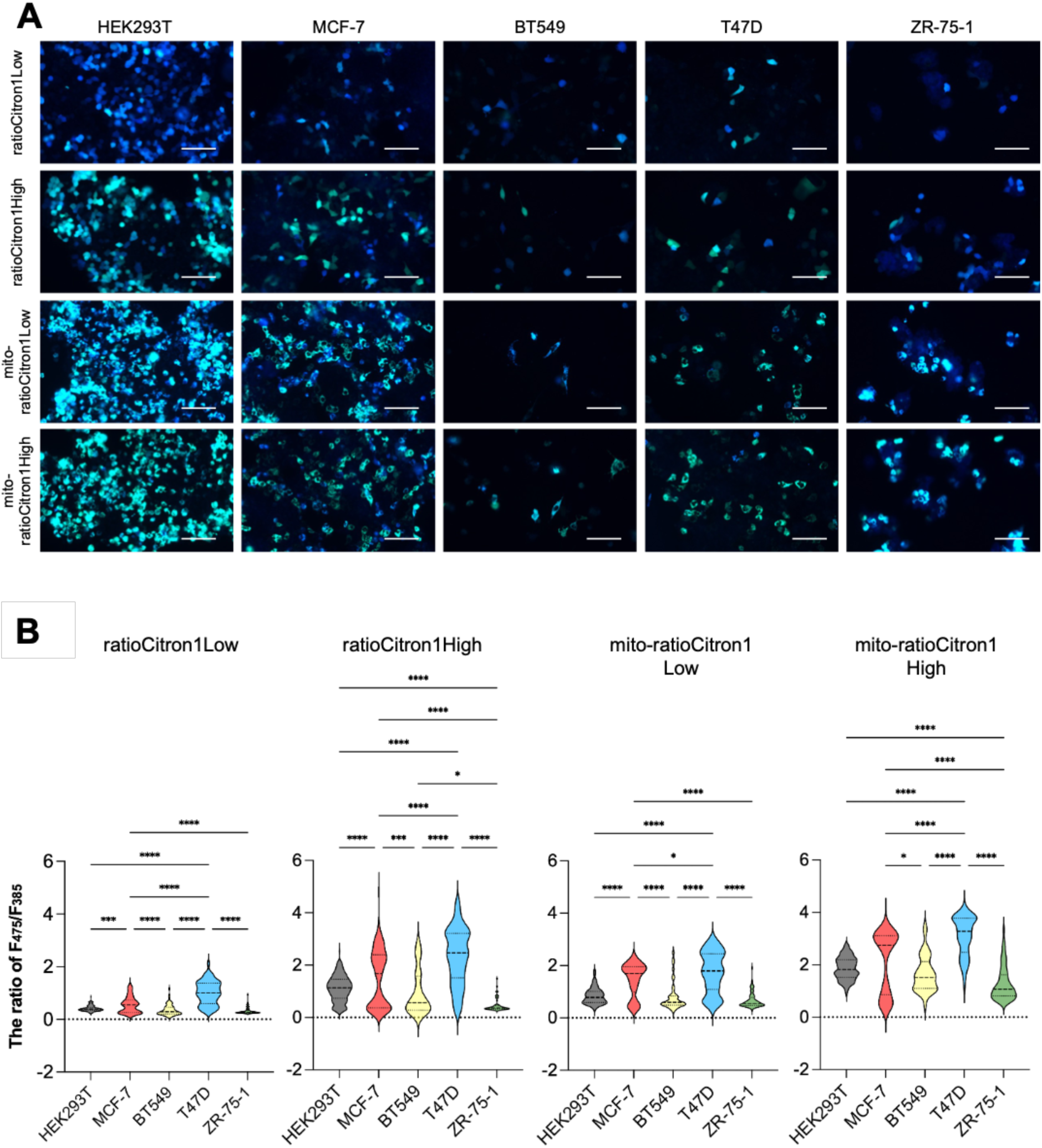
RatioCitron1 uncovers the heterogeneity of citrate levels in human breast cancer cell lines. **A.** Human breast cancer cells and HEK293T cells were transfected with ratioCitron1 variants expressed in cytosol and mitochondria. Representative images are shown. The blue color showed the fluorescent signals taken by 385 nm excitation (emission 499-529 nm) and green color showed the fluorescent signals taken by the 475 nm excitation (emission 500-550 nm). The blue and green overlaid images are shown. Scale bar: 100 µm. **B.** The emission ratio of each excitation wavelength (F_475_/F_385_) is shown (ratioCitron1Low; HEK293T *n* = 160, MCF-7 *n* = 77, BT549 *n* = 64, T47D *n* = 58, ZR-75-1 *n* = 56, ratioCitron1High; HEK293T *n* = 186, MCF-7 *n* = 137, BT549 *n* = 45, T47D *n* = 56, ZR-75-1 *n* = 56, mito-ratioCitron1Low; HEK293T *n* = 127, MCF-7 *n* = 100, BT549 *n* = 43, T47D *n* = 61, ZR-75-1 *n* = 65, mito-ratioCitron1High; HEK293T *n* = 131, MCF-7 *n* = 84, BT549 *n* = 37, T47D *n* = 66, ZR-75-1 *n* = 62). This experiment was independently repeated three times and one result from the representative experiment is shown. Statistically analyzed using one-way ANOVA with Tukey’s multiple comparisons test (*, *p* < 0.05; **, *p* < 0.01; ***, *p* < 0.001; ****, *p* < 0.0001).

### Mitochondria localized ratioCitron1 uncovers metabolic heterogeneity in mouse embryonic stem cells established by differential expression of electron transport chain complexes

Next, we adapted this methodology for flow cytometry–based ratiometric analysis of subcellular metabolite pools. This enables us to make quantitative, population-level measurements of citrate levels with single-cell resolution. mESCs provide a well-defined system for this approach, as they exhibit pronounced metabolic plasticity and intrinsic cell-to-cell heterogeneity while remaining amenable to genetic engineering^23^. We therefore generated a panel of ratioCitron1 knock-in (KI) mESC lines that express either mitochondrial- or cytosol-targeted biosensors with distinct citrate affinities.

The mitochondrial ratioCitron1Low and ratioCitron1High KI cell lines, as well as the cytosolic ratioCitron1High, were successfully established. However, the cytosolic ratioCitron1Low KI line did not yield a detectable fluorescent signal (data not shown). We hypothesised that this reflected the relatively low citrate abundance in the cytosol, similar to what we observed in the cancer cells (**Fig. 4B**). Consistent with this interpretation, metabolomic analysis of biochemically fractionated samples revealed that cytosolic citrate concentrations are substantially lower than mitochondrial concentrations (**Supplementary Fig. 6F‒G**), indicating that only the ratioCitron1High biosensor enables reliable detection of cytosolic citrate in stably expressing mESCs.

Imaging flow cytometry further confirmed that both mitochondrial ratioCitron1 variants colocalised with MitoTracker-labelled mitochondria, whereas the cytosolic ratioCitron1High did not (**Supplementary Fig. 6B‒C**). This validates correct subcellular targeting and establishes the suitability of these KI lines for compartment-specific citrate measurements by flow cytometry.

To determine whether the biosensors report physiologically relevant changes in citrate metabolism, we perturbed the TCA cycle by siRNA-mediated depletion of citrate synthase (Cs) or mitochondrial aconitase 2 (Aco2) (**Supplementary Fig. 6A**). In mitochondria, Cs catalyses citrate formation from acetyl-CoA and oxaloacetate, whereas Aco2 converts citrate to isocitrate. Consistent with these roles, Cs depletion led to a reduction^24^, and Aco2 depletion to an increase^25^, in the mito-ratioCitron1Low signal (**Fig. 5A**, **Supplementary Fig. 6D**). Comparable responses were observed in mito-ratioCitron1High KI mESCs (**Supplementary Fig. 6E**). These results indicate that mitochondrial ratioCitron1 KI cell lines respond to perturbations of citrate production and consumption in the expected direction. Because citrate is exported from mitochondria to the cytosol, changes in mitochondrial citrate availability were also reflected in the cytosolic compartment. Accordingly, Cs depletion decreased, whereas Aco2 depletion increased, the ratio of cytosolic ratioCitron1High (**Fig. 5B**), indicating coordinated regulation of citrate pools across compartments.

**Figure 5.**
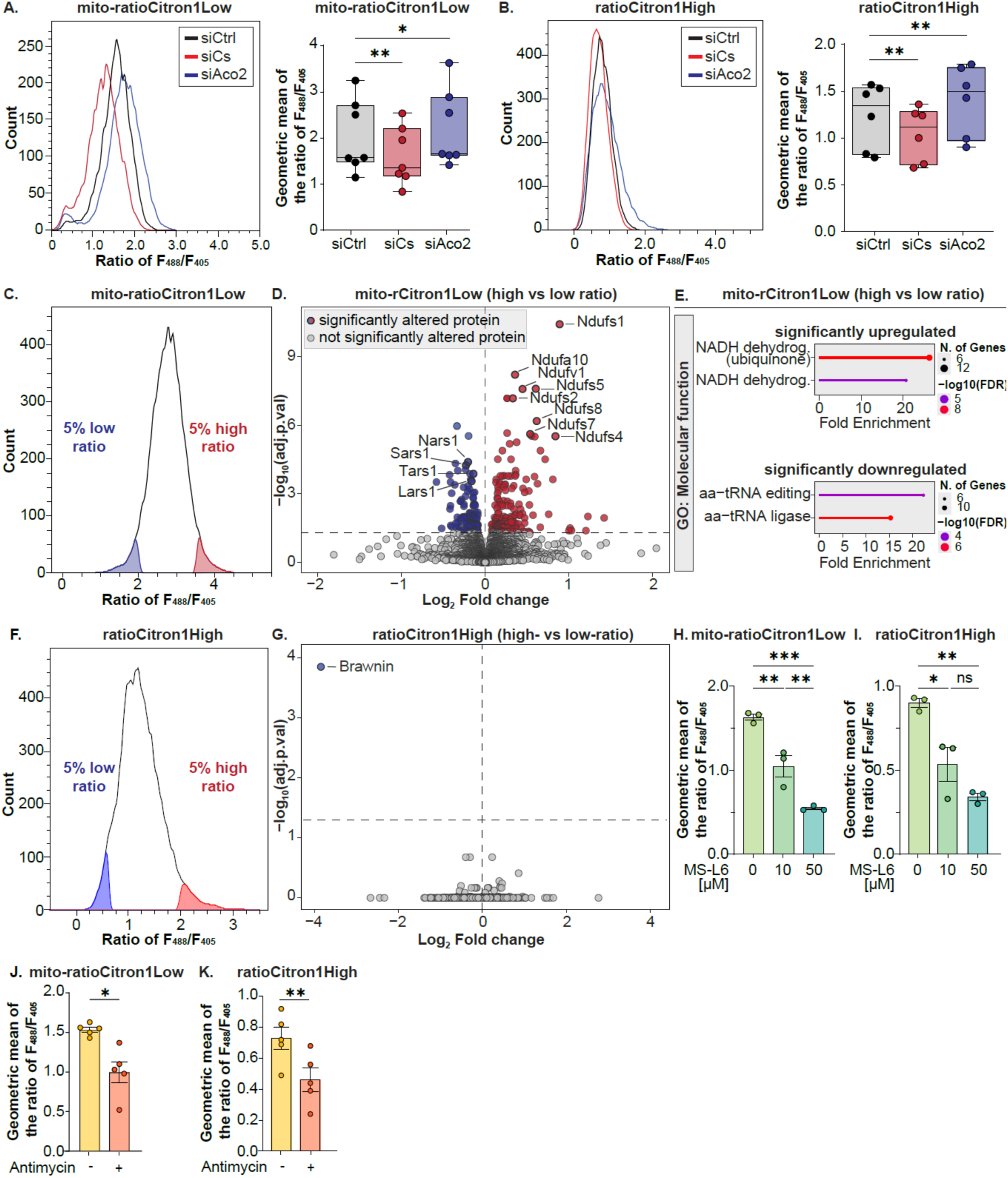
Mitochondrially localized ratioCitron1 uncovers metabolic heterogeneity in mouse embryonic stem cells established by differential expression of electron transport chain complexes. **A–B.** mito-ratioCitron1Low (**A**) or ratioCitron1High (**B**) knock-in (KI) mESCs were treated with non-targeting (siCtrl), citrate synthase (siCs)- and aconitase 2 (siAco2)-targeting siRNAs, respectively, and analyzed by flow cytometry. Representative histograms and the geometric mean of the F_488_/F_405_ ratio is shown. siCtrl, siCs and siAco2-treated cells are highlighted in gray, red and blue, respectively. *n* = 6–7 biological replicates. The box plot represents the median and 25–75 percentiles. Statistically analyzed using one-way ANOVA with Tukey’s multiple comparisons test (*, *p* < 0.05; **, *p* < 0.01). **C.** Representative histogram depicting the strategy selecting the 5% of high-ratio (red) and 5% of low-ratio (blue) cell populations. **D.** Volcano plot depicting the differentially expressed proteins in the high- and low-ratio mito-ratioCitron1 KI cell populations. Significantly up- (red) and downregulated (blue) proteins are highlighted. Selected proteins related to the electron transport chain (ETC) in the upregulated and aminoacyl-tRNA synthases in the downregulated fraction are labelled. **E.** Gene ontology analysis of significantly up- or downregulated proteins as shown in (**D**). **F–G.** Same setup as in (**C–D**) for the ratioCitron1High KI mESCs. **H–K.** mito-ratioCitron1Low (**H and J**) or ratioCitron1High (**I and K**) KI mESCs were cultured in the absence or presence of 10 or 50 µM of the ETC-I inhibitor, MS-L6 for 1 h (**H–I**), or 5 nM ETC-III inhibitor, Antimycin for 2 h (**J–K**) and analyzed by flow cytometry. Geometric mean of the F_488_/F_405_ ratio is shown and statistically analyzed using Student’s *t*-test or one-way ANOVA with Tukey’s multiple comparisons test (*, *p* < 0.05; **, *p* < 0.01; ***, *p* < 0.001). *n* = 3 (**H–I**) and *n* = 5 (**J–K**) biological replicates. Error bars indicate the SEM.

Although the ratioCitron1High biosensor could effectively measure citrate in the mitochondria, its dynamic range appeared more restricted than that of the ratioCitron1Low variant in the mitochondria (**Fig. 5A‒B**, **Supplementary Fig. 6D**), emphasizing the importance of matching biosensor affinity to the physiological concentration range of the target compartment.

Flow cytometric analysis revealed pronounced cell-to-cell heterogeneity in citrate levels between KI mESC populations. To identify molecular features associated with distinct citrate states, we isolated cells representing the top and bottom 5% of the ratiometric distribution and performed proteomic profiling (**Fig. 5C and 5F, Supplementary Table 1**). In the high-citrate population of mito-ratioCitron1Low mESCs, proteins of the electron transport chain (ETC) were significantly enriched (**Fig. 5C‒E**). This trend was also observed in the high-citrate population of mito-ratioCitron1High KI mESCs (**Supplementary Fig. 6H‒I**). The TCA cycle and the ETC are metabolically interdependent: the oxidation of NADH and FADH₂ by ETC complexes sustains TCA cycle flux, while TCA-derived reducing equivalents support oxidative phosphorylation (OXPHOS)^26^. These data suggest that mitochondrial citrate levels correlate with both the ETC activity and the abundance of respiratory chain components, indicating coordination between metabolic state and mitochondrial protein expression.

In contrast, in cytosolic ratioCitron1High KI mESCs, the mitochondrial peptide BRAWNIN was the only differentially expressed peptide and was specifically enriched in the low-citrate population (**Fig. 5G**). BRAWNIN, encoded by *C12orf73*, is required for assembly of the ETC complex III^27,28^. BRAWNIN expression is controlled by nutritional stress and negatively associated with AMPK activation^29^. This association suggests that reduced cytosolic citrate may accompany alterations in ETC-III assembly or stability, although the directionality and mechanistic basis of this relationship remain to be established.

To directly test whether ETC activity influences citrate levels, we inhibited ETC complex I (MS-L6^30^) or complex III (antimycin A^31^). Inhibition of either complex resulted in a marked decrease in citrate levels in both mitochondrial and cytosolic compartments (**Fig. 5H‒K**, **Supplementary Fig. 6J‒K**). These findings indicate that ETC function contributes to maintaining intracellular citrate pools, likely by sustaining TCA cycle flux.

In addition to ETC components, tRNA synthetases were enriched in the low-citrate population of mito-ratioCitron1High KI mESCs, suggesting a link between mitochondrial citrate availability and the translational machinery (**Fig. 5C‒E**). Previous studies have shown that acute disruption of the TCA cycle or oxidative stress induces tRNA synthetase expression through the ATF4-dependent integrated stress response (ISR) signalling pathway^32,33^. In addition, nuclear translocation of specific synthetases, such as tyrosyl-tRNA synthetase (TyrRS), can lead to global translational repression^34^. A recent in-depth study showed that citrate accumulation specifically activates the ISR, which is followed by translational repression^35^. The absence of elevated ATF4 expression in the low-citrate population of mito-ratioCitron1High KI mESCs suggests that tRNA synthetase enrichment is not indicative of an ATF4-dependent stress response. Instead, these data suggest a distinct association between low mitochondrial citrate levels and translational regulation under non-stressed conditions. This points to a potential role for citrate as a metabolic signal coordinating mitochondrial function with protein synthesis capacity in pluripotent stem cells.

Collectively, these results establish flow cytometric ratiometric citrate sensing as a robust approach for resolving metabolic heterogeneity in mESCs and reveal a close association between mitochondrial citrate levels, respiratory chain protein expression, and OXPHOS function.

### Increase in citrate levels derived from mitochondria drives productive stem cell differentiation

Stem cell differentiation is accompanied by a metabolic transition from predominantly glycolytic metabolism to increased reliance on the TCA cycle and OXPHOS^36,37^. Removal of the leukaemia inhibitory factor (LIF) induces spontaneous differentiation of mESCs into derivatives of all three germ layers^38,39^. To examine how citrate metabolism changes during stem cell differentiation, we monitored compartment-specific citrate levels using ratioCitron1 KI mESCs.

Consistent with previous reports showing upregulation of TCA cycle enzymes and increased glucose flux into the TCA cycle during differentiation^37^, we observed a progressive increase in mitochondrial citrate and a decrease in cytosolic citrate over seven days following LIF withdrawal (**Fig. 6A‒B**, **Supplementary Fig. 6L**). Given that mitochondrial citrate concentrations exceed cytosolic concentrations (**Supplementary Fig. 6G**), these results suggest a redistribution of citrate metabolism toward the mitochondrial compartment during differentiation.

**Figure 6.**
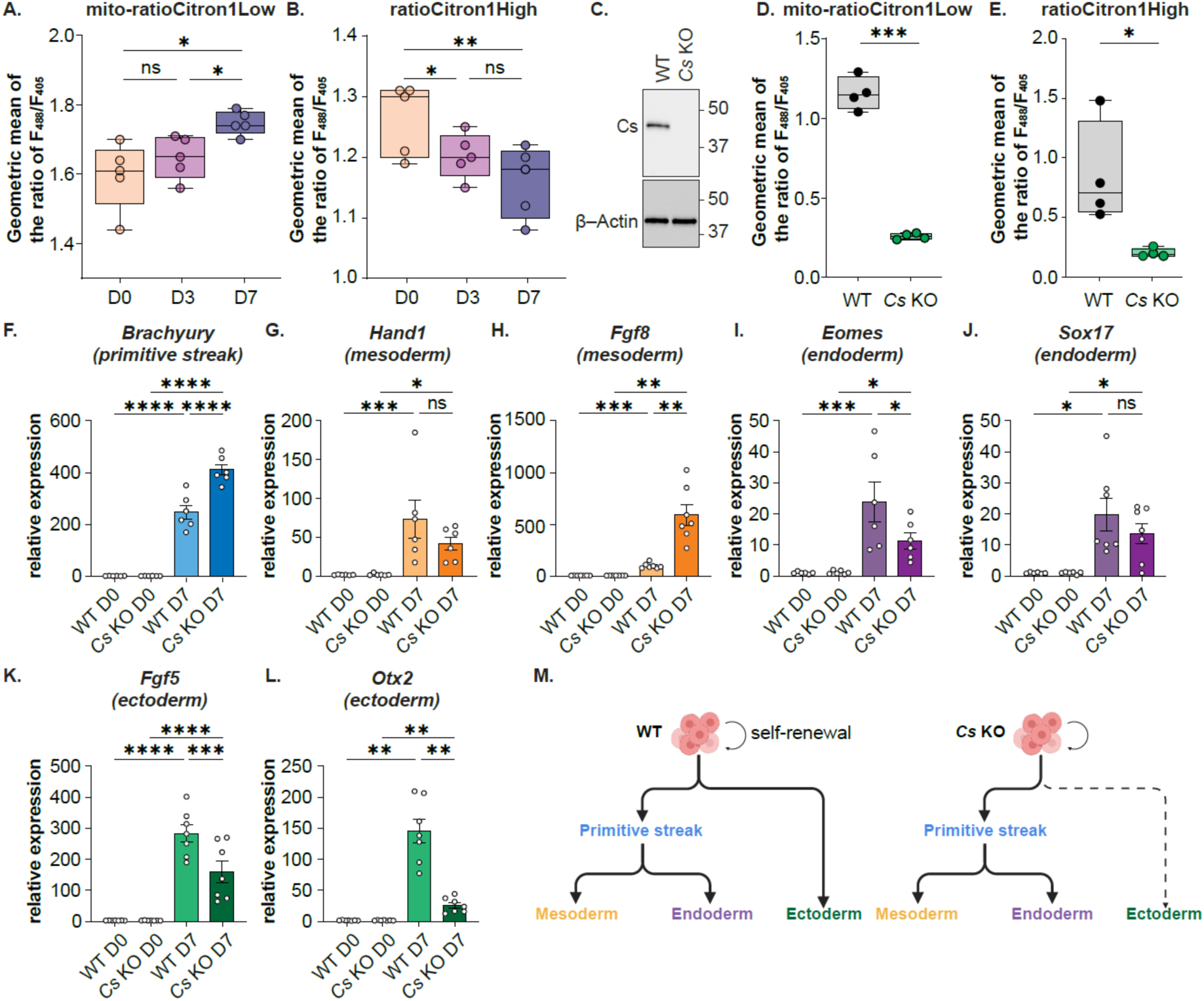
Increase in citrate levels derived from mitochondria drives productive stem cell differentiation. **A–B.** The mito-ratioCitron1Low KI mESCs (**A**) or ratioCitron1High KI mESCs (**B**) were cultured in the presence (D0) or absence of the Leukemia Inhibitory Factor (LIF) for 3 days (D3) or 7 days to induce differentiation (D7). The geometric mean of the F_488_/F_405_ ratio during differentiation is shown and statistically analyzed using or one-way ANOVA with Tukey’s multiple comparisons test (*, *p* < 0.05; **, *p* < 0.01; ns, not significant). *n* = 5 biological replicates. The box plots represent the median and 25–75 percentiles. *n* = 5 biological replicates. **C.** Western blotting of Cs in pluripotent wild-type (WT) and *Cs* KO mESCs was performed. β-Actin was used as a loading control. **D–E.** WT and *Cs* KO mESCs were transiently transfected with the mito-ratioCitron1Low (**D**) or ratioCitron1High (**E**) and the geometric mean of the F_488_/F_405_ ratio is depicted. *n* = 4 biological replicates. Statistically analyzed using or one-way ANOVA with Student’s *t*-test (*, *p* < 0.05; **, *p* < 0.01; ***, *p* < 0.001; ****, *p* < 0.0001; ns, not significant). *n* = 4 biological replicates. The box plots represent the median and 25–75 percentiles. **F–L.** Relative mRNA expression of lineage markers, primitive streak (**F**), mesoderm (**G–H**), endoderm (**I–J**) and ectoderm (**K–L**) in D0 and D7 WT and *Cs* KO mESCs was measured. *n* = 6 biological replicates. Statistically analyzed using one-way ANOVA with Tukey’s multiple comparisons test (*, *p* < 0.05; **, *p* < 0.01; ***, *p* < 0.001; ****, *p* < 0.0001). **M.** The models depicting the trend of stem cell lineage in wild-type and mouse *Cs* KO mESCs during differentiation.

To directly assess the role of citrate production, we generated *citrate synthase* knockout (*Cs* KO) mESCs (**Fig. 6C**). Compared with wild-type cells, *Cs* KO cells exhibited significantly reduced ratioCitron1 signals in both mitochondrial and cytosolic compartments (**Fig. 6D‒E**), confirming that Cs is a major contributor to intracellular citrate pools in mESCs. Loss of Cs induced coordinated metabolic and proteomic adaptation. Metabolomic profiling in the pluripotent state revealed reduced steady-state levels of citrate, *cis*-aconitate, isocitrate, α-ketoglutarate, glutamate, and glutamine. This was accompanied by increased levels of succinate, fumarate, malate, aspartate, and asparagine (**Supplementary Fig. 7E and Supplementary Table 2**), consistent with the diversion of oxaloacetate towards amino acid biosynthesis when citrate production is impaired^24^. Proteomic analysis revealed downregulation of ETC components and upregulation of aminoacyl-tRNA synthetases (**Supplementary Fig. 7A‒B and Supplementary Table 3**), mirroring the enrichment of tRNA synthetases observed in the low-citrate population of mito-ratioCitron1High KI mESCs identified by ratiometric sorting (**Fig. 5C‒E**, **Supplementary Fig. 6H‒L and Supplementary Table 1**). Notably, this remodelling again occurred without activation of ATF4, indicating a non-stress, metabolic regulatory response rather than a classical integrated stress response. Together, these data support a close association between reduced mitochondrial citrate availability, suppression of ETC protein expression, and compensatory upregulation of the translational machinery.

Finally, we used the *Cs* KO cells to investigate the importance of citrate levels for the productive differentiation of mESCs. After seven days of differentiation, *Cs* KO cells showed reduced levels of the pluripotency markers Sox2 and Stat3 (**Supplementary Fig. 7C‒D**), indicating exit from the pluripotent state. However, lineage specification was altered. *Cs* KO cells exhibited increased expression of the primitive streak marker *Brachyury* (**Fig. 6F**), suggesting enhanced or prolonged primitive streak–like identity. In contrast, expression of the ectoderm markers *Fgf5* and *Otx2* was significantly reduced (**Fig. 6K‒L**), indicating impaired ectodermal differentiation. Markers of mesoderm and endoderm displayed heterogeneous expression patterns in *Cs* KO cells (**Fig. 6G‒J**), suggesting that loss of citrate synthase disrupts the orderly progression from the primitive streak to later germ layer fates^39,40^. Together, these results indicate that Cs-dependent citrate production is required for efficient ectodermal differentiation and for coordinated execution of downstream lineage programs (**Fig. 6M**).

## Discussion

To decipher the mechanisms of cellular behavior, it is essential to be able to quantitatively detect and evaluate cellular molecules. To achieve this, several types of ratiometric biosensors have been developed. These include excitation ratiometric biosensors for calcium ion (ratiometric-pericam, GEX-GECO, FGCaMP7, Y-GECOs)^41–45^, pH (Ratiometric pHluorin)^46^, hydrogen peroxide (Hyper)^47^ and ATP (ExRai-AKAR, Perceval, QUEEN)^20,48,49^. For metabolites, ratiometric biosensors for lactate (FiLa)^50^ and malonyl-CoA (Malibu)^51^ have been reported. In this work, we developed excitation ratiometric citrate biosensors and characterized their performance as both purified proteins and expressed in mammalian cells.

An important first step that made this work possible was the observation that the T65A mutation leads to an excitation ratiometric response. In some GFP variants, the deprotonated (anionic) state has a peak absorbance at ∼475 nm and exhibits bright green fluorescence, while the protonated (neutral) state has a peak absorbance at ∼400 nm and often exhibits dim fluorescence. However, as long established for some GFP variants, excitation at the shorter wavelength peak (∼400 nm) can produce bright green fluorescence via excited-state proton transfer (ESPT)^52^. ESPT leads to formation of the deprotonated (anionic) excited state from the protonated (neutral) excited state^52,53^. In the absence of citrate, ratioCitron1 variants are primarily in the protonated state and exhibit bright emission when excited at ∼400 nm due to high ESPT efficiency. When bound to citrate, the ground state equilibrium shifts towards the anionic state, decreasing the emission intensity when excited at ∼400 nm and increasing the emission intensity when excited at ∼495 nm. We expect that the T65A mutation has altered the hydrogen bond network in the vicinity of the chromophore, changing the equilibrium between the protonated and deprotonated states of the ground state chromophore. This equilibrium is further altered by the conformational change in the CitA domain that is communicated to the chromophore environment via the connecting linkers. However, the detailed mechanism remains unclear, and further research may be required to address this question.

With the aim of developing non-responsive control biosensors, previous studies have introduced mutations into the binding pocket to disrupt analyte binding. This approach diminishes the conformational change of the SD and thereby abolishes the fluorescence response^17,19,54^. Although this strategy is often effective, it does not always completely eliminate biosensor responsiveness. In addition, because these control biosensors cannot bind the analyte, the intracellular concentration of free analyte may differ between cells expressing the functional biosensor and those expressing the control variant. The strategy presented in this study instead involves inserting a single glycine residue into the linker region. This modification disrupts transmission of the SD conformational change to the fluorescent protein (FP), while preserving the analyte binding capability of the SD. As a result, the control biosensor retains analyte binding but lacks a fluorescence response, ensuring comparable intracellular analyte concentrations between experimental and control conditions. This simple linker insertion provides an effective and generalizable method for generating non-responsive control biosensors.

By expressing the ratioCitron1 series in multiple breast cancer cell lines, we revealed citrate level heterogeneity at single-cell resolution, which would have been averaged out in other metabolic measurements. This application further demonstrates a key advantage of ratiometric biosensors, in which we can compare the analyte concentrations independent of the biosensor expression levels. Unlike calcium ions, the dynamics of cellular metabolites are relatively moderate and more accurate quantification is essential for biological applications. Breast cancers can be categorized based on the expression of specific receptors, such as the estrogen receptor (ER), progesterone receptor (PR) and human epidermal growth receptor 2 (HER2)^55^. Among the cell lines that we used, the MCF-7, T47D and ZR-75-1 cells are categorized as luminal, which is a less aggressive type^55^. BT549 cells are categorized as triple negative, which is the most aggressive type^55^. The triple negative type shows a higher reliance on glycolysis over the TCA cycle compared to the less aggressive luminal type. These findings underline the clear correlation between the balance of TCA to glycolysis engagement and cancer malignancy^35,56^. In this study, we found higher citrate levels in MCF-7 and T47D cells compared to ZR-75-1 and BT549 (**Fig. 4**), which is partially consistent with malignancy. Based on the biosensor results and metabolomics data (**Supplementary Fig. 7**), we propose higher citrate levels indicate higher TCA cycle activity. Consequently, ZR-75-1 and BT549 cells may rely more readily on glycolysis, a hallmark of cancer malignancy. When we interpret these microscopy data, several features need to be taken into consideration. In this study, all cell lines were subjected to the same conditions, including the seeding cell number, medium used and the microscopy settings (e.g. exposure time and laser power). However, cell density was not controlled for, which could influence metabolism. In addition, the number of the cells within the microscopy fields is limited, so a more comprehensive analysis is required to uncover the relationship between metabolite levels and cancer cell malignancy. Lastly, citrate is in equilibrium within the TCA cycle and controlled by other metabolites such as glutamine, so the TCA activity must be evaluated carefully.

It is technically challenging to use metabolomic analyses to distinguish citrate originating from different compartments, and it is difficult to monitor citrate levels in live cells. In contrast, the ratioCitron1 biosensors can be specifically targeted to mitochondria or can be expressed in the cytosol. In this study, we employed compartment-specific monitoring of citrate changes using the ratioCitron1 during differentiation. This revealed an increase in mitochondrial citrate and a decrease in cytosolic citrate (**Fig. 6A**). Mitochondrial citrate is essential for the TCA cycle and OXPHOS to produce ATP^37^, indicating that increased mitochondrial citrate primarily supports energy production. Conversely, cytosolic citrate levels decline during differentiation. Cytosolic citrate is a key source of acetyl-CoA, which contributes to *de novo* lipid synthesis and protein acetylation, including histone acetylation, involved in epigenetic regulation^57^. Previous studies have shown that histone lysine acetylation decreases during early stages of differentiation, contributing to chromating remodelling. In addition, *de novo* fatty acid synthesis is downregulated during differentiation^58^, and lipid composition undergoes significant changes^59^, supporting a shift in citrate-dependent metabolic outputs under conditions of low cytosolic citrate. Taken together, these findings suggest that citrate partitioning shifts from anabolic to catabolic usage during differentiation. This approach provides a valuable tool to further investigate how limited citrate availability coordinates epigenetic regulation and lipid metabolism in differentiated cells.

Towards this goal, we generated *Cs* knock-out mESCs. *Cs* knockout reduces intracellular citrate availability, leading to global metabolic rewiring and altered lineage specification. Reduced citrate limits cytosolic acetyl-CoA production, which is predicted to alter histone and global protein acetylation and thereby reprogram transcriptional states required for proper differentiation. Changes in tRNA synthetases further suggest coupling of metabolic status to translational control. Concomitant alterations in tRNA synthetase expression indicate that the citrate perturbation extends to translation (**Supplementary Fig. 7A‒B**). Collectively, these observations support a coordinated axis linking metabolism to epigenetic regulation and translation, ultimately influencing cell fate determination.

## Materials and Methods

### General methods and materials

All DNA primers were purchased from Integrated DNA Technologies (IDT) or Thermo Fisher Scientific. pBAD-Citron1 (Addgene Plasmid #134300) was used as a template. Phusion high-fidelity DNA polymerase (Thermo Fisher Scientific) and CloneAmp^TM^ HiFi PCR Premix (TaKaRa, 639298) were used for routine polymerase chain reaction (PCR) amplification, and Taq DNA polymerase (New England Biolabs) was used for error-prone PCR. A QuickChange mutagenesis kit (Agilent Technologies) was used for site-directed mutagenesis. Restriction endonucleases and rapid DNA ligation kits (Thermo Fisher Scientific) were used for plasmid construction. Citric acid (CALEDON, 3020-1) or trisodium citrate dihydrate (Wako, 204-16675) was used for the assays. Products of PCR and restriction digests were purified using agarose gel electrophoresis and the GeneJET gel extraction kit (Thermo Fisher Scientific, K0691). Plasmids were extracted from *E. coli* using the GeneJet Plasmid Miniprep Kit (Thermo Fisher Scientific, K503) and proteins were extracted by B-PER (bacterial protein extraction reagent, Thermo Fisher Scientific, 78248). DNA sequences were analyzed by the DNA sequence service of Fasmac Co. Ltd or University of Alberta Molecular Biology Services Units. The fluorescence spectra and intensity were recorded on Safire2 (Tecan) or Spark plate reader (Tecan). For ETC inhibitor treatment of mESCs, MS-L6 (MedChemExpress, HY-158421) and Antimycin A (Sigma-Aldrich, A8674) were used.

### Structural modeling of ratioCitron1

The modeling structures of ratioCitron1 were generated by AlphaFold^21^. The PyMOL (https://www.pymol.org) was used for structural analysis.

### Engineering of ratioCitron1 variants

pBAD-Citron1 was used as a template and the T65A point mutation was inserted. Site directed mutagenesis was performed using CloneAmp^TM^ HiFi PCR Premix (TaKaRa, 639298). For protein expression, plasmids were expressed in *E.coli* DH10B and cultured in LB medium supplemented with 100 µg/mL ampicillin (Fisher, BP1760-25) and 0.02% w/v L-arabinose (Alfa aesar, A11921). The *E.coli* was grown at 37°C overnight and proteins were extracted by B-PER on the following day. Directed evolution with random mutagenesis was performed as previously described^17^. Briefly, the error-prone PCR (ER-PCR) was conducted with Taq polymerase (New England Biolabs). PCR products and pBAD vectors digested by restriction enzymes (New England Biolabs or Thermo Fisher Scientific) were purified by a GeneJET gel extraction kit. The purified PCR fragments were inserted into pBAD plasmid by In-Fusion Snap Assembly Master Mix (TaKaRa, 638948).

### Protein purification

The ratioCitron1 variants in pBAD expression vectors containing a N-terminal poly His (6×) tag were expressed in *E. coli* BL21 (DE3). Colonies were used to inoculate LB supplemented with 100 μg mL^−1^ ampicillin and grown at 37°C until an OD600 value of 0.6 was reached. The cell culture was then induced by adding 0.04% (w/v) L-arabinose and grown overnight at room temperature. The next day, cells were harvested by centrifugation. The cell pellet was resuspended in lysis buffer (50 mM Na_2_HPO_4_, 300 mM NaCl, 10 mM imidazole, pH 8.0) and B-PER. Cells were placed on ice and lysed by sonication (50 % on/off for 2 min) and then centrifuged. Lysates were filtered with a 0.45 μm filter and loaded onto a lysis buffer pre-equilibrated Ni-NTA column. The column was then washed with 15 column volumes (cv) of wash buffer (50 mM Na_2_HPO_4_, 300 mM NaCl, 250 mM imidazole, pH 8.0) and buffer exchanged into citrate (−) buffer (TBS, pH 7.2) using a centrifugal spin column (10K MWCO, Thermo Fisher Scientific).

### *In vitro* characterization

Purified proteins were used for all *in vitro* characterizations. All the fluorescence spectra were collected by a plate reader (TECAN) using Greiner 96/384-well flat-bottom microplates. For citrate titration, buffers were prepared by mixing citrate (−) buffer (TBS, pH 7.2) and citrate (+) buffer (1 M trisodium citrate dihydrate, pH 7.2) to provide citrate concentrations ranging from 0 mM to 1 M. To perform pH titrations, 10 µL protein solutions were diluted into 90 µL buffers that contained 100 mM NaCl, 30 mM sodium acetate, 30 mM MOPS, 30 mM Tris, and 30 mM sodium tricarbonate and either no citrate or 10 mM trisodium citrate dihydrate. The solution was adjusted to pH values ranging from 4 to 11. Absorbance spectra were measured using the Shimadzu UV1800 spectrometer.

### HeLa cell imaging

For mammalian cell imaging, a series of ratioCitron1 biosensors were transferred to pcDNA3. HeLa cells [American Type Culture Collection (ATCC)] were maintained in Dulbecco’s modified Eagle’s medium (DMEM high glucose; Nacalai Tesque, 08459-35) supplemented with 10% fetal bovine serum (FBS; Sigma-Aldrich) and 100 μg mL^−1^ penicillin/streptomycin (Nacalai Tesque, 09367-34). Cells were transiently transfected with the plasmids with Lipofectamine 3000 (Thermo Fisher) in Opti-MEM I (Gibco, 31985062) and imaged within 24-72 h after transfection. Cells were seeded on a 35 mm/glass base dish (Glass 12 Φ, IWAKI, 3911-035, Glass 27 Φ, IWAKI, 3910-035). An IX83 wide-field fluorescence microscope (Olympus) equipped with a pE-300 LED light source (CoolLED) and a 40× objective lens (numerical aperture (NA) = 1.3; oil) or a 20× objective lens (NA = 0.8; air), an ImagEM X2 EM-CCD camera (Hamamatsu), and Cellsens software (Olympus) was used for imaging. The filter sets for imaging had the following specifications. ratioCitron1 variants: 405/20 nm, dichroic mirror 425 nm dclp, and emission 518/45 nm and excitation 470/20 nm, dichroic mirror 490 nm dclp, and emission 518/45 nm. Fluorescent images were analyzed with ImageJ software (https://imagej.net/software/fiji/, National Institutes of Health; accessed on September 1, 2023)^60^. For imaging the treatment with MPC inhibitor UK-5099 (MedChemExpress) and ACLY inhibitor BMS-303141 (MedChemExpress), Hank’s balanced salt solution (HBSS; Nacalai Tesque, 09735-75) and 10 mM HEPES (Nacalai Tesque, 177557-94) was used as imaging buffer. Each reagent was stored in 10 mM in DMSO and prepared to be desired concentration diluted with an imaging buffer.

### Cell imaging of cancer cell lines

HEK293T (ATCC) cells were cultured in DMEM high glucose (Gibco, 11965118) with 10% FBS (Gibco) and penicillin/streptomycin (Gibco, 15140122). BT549 (ATCC), T47D (ATCC) and ZR-75-1 (ATCC) cells were cultured in RPMI1640 (Gibco, 11965092) supplied with 10% FBS and penicillin/streptomycin. MCF-7 (ATCC) cells were cultured in MEM (Gibco) supplied with 10% FBS and penicillin/streptomycin. Cells were seeded on glass bottom plates (Cellvis, D35-20-1.5-N) and after one day incubation, cells were transiently transfected with the ratioCitron variants (pcDNA3) using Lipofectamine 3000 (Thermo Fisher Scientific, L3000015) in Opti-MEM I (Gibco, 31985062). From that time point, all dishes were cultured with RPMI1640 supplied with FBS and penicillin/streptomycin to make all the culturing conditions the same. Two days after transfection, cell imaging was performed. Before taking images, the old medium was removed, washed with HBSS (Ca^2+^,Mg^2+^, without phenol red, Gibco, 14025-092). Then, the images were taken by the Axio Observer Z1/7 (Zeiss) (excitation laser: 385 nm, emission filter: 499-529 nm / excitation laser: 475 nm, emission filter: 500-550 nm). The images were processed with the ZEN software and the quantification analysis was performed with FIJI software. The region of interest (ROI) was set in each cell expressing biosensors and the fluorescent intensity (F) was calculated. The same size of ROI was assigned on the cells without expressing biosensors and the F from that ROI was calculated as background F. The background F was distracted from the F of each targeted ROI. Then, the F from the image taken by the 475 nm laser (F_475_) was divided by the F from the image taken by the 385 nm laser (F_385_) to calculate the ratio of F_475_/F_385_. This experiment was independently repeated three times and one result from the representative experiment is shown.

### Mouse embryonic stem cell culture

AN3-12 mouse embryonic stem cells (mESCs) were cultured as previously described^23^. In brief, DMEM high glucose (Gibco, 41965039) was supplemented with 5 mM of L-glutamine (Sigma-Aldrich, G7513), 15% of FBS (Gibco, A5256801), penicillin/streptomycin (Sigma-Aldrich, P4333), non-essential amino acids (Gibco, 315048), 1 mM of sodium pyruvate (Gibco, 11360039), 0.1 mM of β-mercaptoethanol (Carl ROTH, 4227.3) and 0.02 µg/ml leukemia inhibitory factor (LIF; EMBL Heidelberg) and used to culture cells at 37°C in 5% CO_2_ on tissue culture plates. For induction of spontaneous differentiation of AN3-12 cells, 250 and 1,000 cells were seeded in a 6-well plate and incubated for 3 or 7 days in medium without LIF, respectively. No mycoplasma contamination was detected (Eurofins Genomics).

### Establishment of ratiometric Citron1 Knock-in mESC

For introducing the ratioCitron1variants encoding genes into the *Rosa26* locus, we used the protocol and constructs as previously described^61^. The individual ratioCitron1 encoding sequence was amplified by PCR and introduced into the donor plasmid for knock-in (KI) using restriction-free cloning. Haploid mESCs were transfected with the Cas9-mRFP and *Rosa26* targeting guide RNA encoding plasmid and the donor plasmid using Lipofectamine 3000 (Invitrogen, L3000008) according to the manufacturer’s protocol. GFP and mRFP-positive cells were sorted by the BD FACSAria Fusion cell sorter and cultured for several days. After single cell sorting into 96 well plates, clones were analyzed for transgene insertion by genotyping PCR and the intensity of ratioCitron1 was confirmed by flow cytometry (BD FACSCantoII). To sort diploid mESCs for further analysis, cells were stained with Hoechst 33342 (Enzo Biochem, ENZ-52401). To exclude dead cells, propidium iodide (AppliChem, A2261) staining was added. Cells were sorted for DNA content on a BD FACSAria Fusion sorter and flow profiles were recorded with the FACSDiva software (BD Franklin Lakes). All subsequent experiments were conducted in diploid mESCs. Primers for genotyping PCR are listed in **Supplementary Table 4**.

### Establishment of *Citrate synthase* Knock-out mESC

CRISPR guide RNA sequence, which was designed for mouse *Citrate synthase* (*Cs*) gene, was cloned into a Cas9-mRFP encoding vector. The target sequence was 5′-GCAGCAACATGGGAAGACAG-3′. After 48 h-post transfection, mRFP-positive cells were isolated using a BD FACSAria Fusion cell sorter. A haploid clone with a frame shift mutation in *Cs* allele was identified by genomic PCR and DNA sequencing. Primers for Genomic PCR are listed in **Supplementary Table 4**.

### Small interfering RNA (siRNA) treatment

SMARTpools of ON-TARGETplus siRNAs (Dharmacon, Lafayette, CO) targeting mouse *Citrate synthase* (L-059581), *Aconitase 2 mitochondrial* (L-065502) or non-targeting control siRNA (D-001810) were used. siRNAs were transfected using Lipofectamine RNAiMAX (Invitrogen, 13778-150) according to the manufacturer’s protocol. In brief, 20 nM of 1 µl of each siRNA pool and 3 µl Lipofectamine RNAiMAX were mixed in Opti-MEM I medium (Gibco, 31985062) and incubated at room temperature for 15 minutes. Cells were incubated for 48 h after transfection to allow for efficient knockdown before conducting further experiments.

### Flow cytometry analysis

The cells were harvested with 0.1% Trypsin/EDTA (Gibco, 15400054). Harvested cells were re-suspended in flow cytometry buffer supplemented with 15 % FBS, HBSS, penicillin/streptomycin, non-essential amino acids, sodium pyruvate, β-mercaptoethanol and LIF and analyzed by flow cytometry (BD FACSCantoII). Dead cells were excluded by DRAQ7 (BioLegend, 424001) staining. mESCs transiently transfected with plasmids for mito-Citron1^19^ and mT-Sapphire were used for compensation in all experiments. mT-Sapphire was cloned from the Peredox vector (Addgene; 32383)^62^. The same bandpass filter, 525/50 to detect both 488 laser- and 405 laser-excited signals was equipped in FACSCanto-II. For flow cytometry analysis using ratioCitron1 KI mESC lines, we gated and analyzed only the biosensor positive positive population. Flow data was recorded with the FACSDiva software (BD) and analyzed with FlowJo 10 software (BD). For Imaging flow cytometry, the cells were stained with MitoTracker Red CMXRos (Invitrogen, M46752) to visualize mitochondria for 20 min before harvesting. The bandpass filters, 528/65 to detect 488 laser- and 537/65 to detect 405 laser-excited signals were equipped in Amnis ImageStreamX Mk II. Cell images and flow data was recorded using the IDEAS software and analyzed with the INSPIRE software. Fluorescence intensity profiles were analyzed by Fiji software.

### Western Blotting

Cells were lysed with ice cold Triton-X-based lysis buffer (1% Triton-X, 50 mM Tris-HCl (pH 7.5), 150 mM NaCl and 1 mM EDTA), supplemented with Protease inhibitor cocktail (Roch, 11873580001). After removing cell debris by centrifugation at 11,000×*g* for 10 min, protein concentration of cell lysates was determined using the Pierce^TM^ BCA protein assay kit according to manufacturer’s instructions (Thermo Scientific, 23227). Samples were adjusted in sodium dodecyl sulfate (SDS) based sample buffers (2% SDS, 50 mM Tris-HCl (pH 6.8), 10 % glycerol, 0.1 M dithiothreitol and 0.01% Bromophenol blue). After boiling the samples at 70°C, equal protein amounts were subjected to SDS-PAGE and transferred to a nitrocellulose membrane using the Trans-Blot Turbo Transfer system (BioRad). After blocking with 3% skim milk (SeraCare, 42590.02) for 30 min, membranes were incubated with specific antibodies overnight at 4°C. All antibodies were used in 1% skim milk in 0.01% TBS-T. After incubation with HRP-conjugated secondary antibodies, the blot was developed using ECL solution (Merck Millipore, WBKLS0500) on a ChemiDoc MP Imaging System (BioRad). The following antibodies were used as primary antibodies in this study: Aconitase 2 (Proteintech, 67509-1-Ig, 1:5,000, RRID:AB_2882730), Citrate Synthase (Cell Signaling Technology, 14309S, 1:5,000, AB_2665545), HSP90 (Cell Signaling Technology, 4874, 1:10,000, RRID:RRID:AB_2121214), SDHA (Thermo Scientific, 459200, 1:5,000, RRID:AB_10838019), SOX2 (ABclonal, A19118, 1:8,000, RRID:AB_2862611), Stat3 (Cell Signaling Technology, 9139, 1:5000, RRID:AB_331757), TOMM20 (Sigma-Aldrich, HPA011562, 1:5,000, RRID:AB_1080326) Vinculin (ABclonal, A2752, 1:10,000, RRID:AB_2863020) β-Actin (Sigma-Aldrich, A5441, 1:15,000, RRID:AB_476744). The following antibodies were used as secondary antibodies rabbit IgG (Thermo Fisher Scientific, AB2536530, 1:15,000), and mouse IgG (Thermo Fisher Scientific, AB2536527, 1:15,000).

### Cell Fractionation

AN3-12 cells were seeded in five 10 cm dishes at a density of 0.5 × 10^6^ cells per dish as five technical replicates and cultured for 48 h. Cells were harvested and resuspended in the fractionation buffer (10 mM Tris-HCl pH 7.5, 10 mM KCl and 1.5 m MgCl_2_) containing protease inhibitors. The whole cell was taken. Samples were kept on ice for 20 min and subsequently processed by syringing with a 27G needle. Cell debris and nuclei fraction were removed by centrifuging the cell homogenate at 1,000×*g* for 10 min. The supernatant was centrifuged at 11,000×*g* for 10 min to separate the cytosol. Crude mitochondria fraction was isolated at 11,000×*g* for 10 min and washed twice with the fractionation buffer. All processes were performed at 4°C. The fractions were processed for downstream applications. Five percent of fractionated lysate was taken for protein quantification with Pierce BCA protein assay kit and the measured protein amounts were used for metabolite normalization.

### Anion exchange chromatography mass spectrometry (IC-MS) analysis

The cells were washed twice, using 1 mL of 75 mM ammonium carbonate pH 7.4 (Sigma) wash buffer warmed to 37°C. Metabolites were extracted by adding 400 µL of pre-cooled (-20°C) metabolite extraction buffer (40:40:20 (v:v:v) acetonitrile:methanol:water (Optima LC/MS grade, Thermo Fisher Scientific)). The whole lysate, including the precipitated cellular material, was transferred to a new tube and stored on ice. The lysates were centrifuged for 10 min at 21,000×*g* at 4°C. The supernatant was transferred to a new tube and the polar metabolite extract was dried immediately in a SpeedVac concentrator (ScanVac) with the following settings (20°C at 1,000 rpm). Samples were measured on the diverse LC-MS systems.

Semi-targeted liquid chromatography-high-resolution mass spectrometry-based (LC-HRMS) analysis of amine-containing compounds was performed using an adapted benzoylchlorid-based derivatization method^63^. In brief: The polar fraction of the metabolite extract was re-suspended in 100 µL of LC-MS-grade water (Optima-Grade, Thermo Fisher Scientific) and incubated at 4°C for 15 min on a thermomixer. The re-suspended extract was centrifuged for 5 min at 21,100×*g* at 4°C and 25 µL of the cleared supernatant were mixed in an auto-sampler vial with a 300 µL glass insert (Chromatography Accessories Trott, Germany). The aqueous extract was mixed with 12.5 µL of 100 mM sodium carbonate (Sigma), followed by the addition of 12.5 µL 2% [v/v] benzoyl chloride (Sigma) in acetonitrile (Optima-Grade, Thermo Fisher Scientific). Samples were vortexed and kept at 20°C until analysis.

For the LC-HRMS analysis, 2 µL of the derivatized sample was injected onto a 100 x 2.1 mm Premier HSS T3 UPLC column (Waters). The flow rate was set to 400 µL/min using a binary buffer system consisting of buffer A (10 mM ammonium formate (Sigma), 0.15% [v/v] formic acid (Sigma) in LC-MS-grade water. Buffer B consisted of acetonitrile (Optima-grade, Thermo Fisher-Scientific) containing 0.1% formic acid (Sigma). The column temperature was set to 40°C and the LC gradient was: 0% B at 0 min, 0–15% B 0–4.1min; 15–17% B 4.1–4.5 min; 17–55% B 4.5–11 min; 55–70% B 11–11.5 min, 70–100% B 11.5 –13 min; B 100% 13–14 min; 100–0% B 14–14.1 min; 0% B 14.1–19 min; 0% B. The mass spectrometer (Orbitrap Plus, Thermo Fisher Scientific) was operating in positive ionization mode recording the mass range m/z 100–1000. The heated ESI source settings of the mass spectrometer were: Spray voltage 3.5 kV, capillary temperature 300°C, sheath gas flow 60 AU, aux gas flow 20 AU at a temperature of 340°C and the sweep gas to 2 AU. The RF-lens was set to a value of 60%. The semi-targeted data analysis for the samples was performed using the TraceFinder software (Version 5.1, Thermo Fisher Scientific). The identity of each compound was validated by authentic reference compounds, which were run before and after every sequence. Peak areas of [M + nBz + H]^+^ ions were extracted using a mass accuracy (<5 ppm) and a retention time tolerance of <0.05 min. Areas of the cellular pool sizes were normalized to the internal standards (U-^15^N; U-^13^C amino acid mix (MSK-A2-1.2), Cambridge Isotope Laboratories), which were added to the extraction buffer, followed by a normalization to the fresh weight of the analyzed sample.

For Anion-Exchange Chromatography Mass Spectrometry (AEX-MS) for the analysis of anionic metabolites, 75 µL of the cleared supernatant described in the section for the analysis of amine-containing polar metabolites, were transferred to polypropylene autosampler vials (Chromatography Accessories Trott, Germany). The samples were analyzed using a Dionex ionchromatography system (Integrion, Thermo Fisher Scientific) as described previously^64^. In brief, 5 µL of polar metabolite extract were injected in push partial mode using an overfill factor of 1, onto a Dionex IonPac AS11-HC column (2 mm × 250 mm, 4 μm particle size, Thermo Fisher Scientific) equipped with a Dionex IonPac AG11-HC guard column (2 mm × 50 mm, 4 μm, Thermo Fisher Scientific). The column temperature was held at 30°C, while the auto sampler was set to 6°C. A potassium hydroxide gradient was generated using a potassium hydroxide cartridge (Eluent Generator, Thermo Scientific), which was supplied with deionized water. The metabolite separation was carried at a flow rate of 380 µL/min, applying the following gradient conditions: 0–3 min, 10 mM KOH; 3–12 min, 10–50 mM KOH; 12–19 min, 50–100 mM KOH, 19–22 min, 100 mM KOH, 22–23 min, 100–10 mM KOH. The column was re-equilibrated at 10 mM for 3 min before injecting the next sample. For the analysis of metabolic pool sizes the eluting compounds were detected in negative ion mode using full scan measurements in the mass range m/z 50–750 on a Q-Exactive HF high resolution MS (Thermo Fisher Scientific). The heated electrospray ionization (ESI) source settings of the mass spectrometer were: Spray voltage 3.2 kV, capillary temperature was set to 300°C, sheath gas flow 60 AU, aux gas flow 20 AU at a temperature of 300°C and a sweep gas glow of 2 AU. The S-lens was set to a value of 60. The semi-targeted LC-MS data analysis was performed using the TraceFinder software. The identity of each compound was validated by authentic reference compounds which were measured at the beginning and the end of the sequence. For data analysis the area of the deprotonated [M-H^+^]^-^ or doubly deprotonated [M-H^2+^]^2-^monoisotopic mass peak of each compound was extracted and integrated using a mass accuracy <5 ppm and a retention time (RT) tolerance of <0.05 min as compared to the independently measured reference compounds. Areas of the cellular pool sizes were normalized to the internal standards, which were added to the extraction buffer, followed by a normalization to the fresh weight of the analyzed sample.

### Whole cell Proteomics analysis

Cell pellets were resuspended in lysis buffer (6 M guanidinium chloride, 2.5 mM tris(2-carboxyethyl) phosphine (TCEP), 10 mM chloroacetamide, 100 mM tris-hydrochloride) and heated to 95°C for 10 min. The lysates were sonicated (30 s/ 30 s, 10 cycles, high performance) using a Bioruptor (B01020001; Diagenode), followed by centrifugation at 21,000×*g* for 20 min. 200 µg of supernatants were digested with 1 µl trypsin (V5280; Promega) overnight at 37°C with shaking. Formic acid was added to the digested peptide lysates (to a final concentration of 1%) to stop trypsin digestion. Samples were desalted by homemade Stage Tips. Eluted lysates in 40% acetonitrile/0.1% formic acid were dried by vacuum centrifugation (Concentrator Plus; Eppendorf) at 45°C. 2 µg of peptide of each sample was dried in a Speed-Vac.

### Cell sorting and lysis for Proteomics analysis

The cells were harvested with 0.1% Trypsin/EDTA and re-suspended in flow cytometry buffer. 5,000 cells from the gated regions were directly sorted into lysis buffer (0.025% n-Dodecyl-β-D-Maltoside,125 mM Triethylammonium bicarbonate buffer, 10 mM TCEP and 20 mM CAA (chloroacetamide)) using FACSAria Illu Svea cell sorter. The lysate was heated to 95°C for 10 min and sonicated (30 s/ 30 s, 10 cycles, high performance) by Bioruptor. Lysate was incubated overnight with a protease mixture (1 µg/µl Trypsin, 0.5 µg/µl LysC in 50 mM acetic acid) at 22°C.

### Liquid chromatography Mass spectrometry

The Evosep One liquid chromatography system^65^ was used for analyzing the samples with the predefined 60 samples per day (60SPD) method. The analytical column we used was an ReproSil-Pur column, 8 cm x 150 µm, with 1.5 µm C18 beads (EV1109 Performance Column, Evosep). The mobile phases A and B were 0.1 % formic acid in water and 0.1% formic acid in 100% ACN, respectively.

Peptides were analyzed on a hybrid TIMS quadrupole TOF mass spectrometer (timsTOF HT, Bruker) in a data-independent acquisition parallel accumulation, serial fragmentation (diaPASEF) mode. The mass spectra range was set to 350 - 1200 m/z and TIMS ion accumulation and ramp times were set to either 50 ms or 100 ms and total cycle time was 0.73 s or 1.38 s. The ion mobility range was set to 1/K0 = 0.8 - 1.25 V-s/cm2. Isolation windows in the m/z versus ion mobility plane were defined to cover the region of highest precursor ion density using py_DIA^66^. Collision energy was applied linearly with ion mobility from 0.6 to 1.6 V-s/cm2, and collision energy from 20 to 59 eV.

### Data Analysis of Proteomics

Raw data was analyzed using Spectronaut version 20.0.25 (Biognosys) using the default parameters against the one-protein-per-gene reference proteome for *Mus musculus*, UP000000589, downloaded August, 2022. Methionine oxidation and protein N-terminal acetylation were set as variable modifications; cysteine carbamidomethylation was set as fixed modification. The digestion parameters were set to “specific” and “Trypsin/P,” with two missed cleavages permitted. Protein groups were filtered for at least two valid values in at least one comparison group and missing values were imputed from a normal distribution with a down-shift of 1.8 and standard deviation of 0.3. Differential expression analysis was performed using limma, version 3.60.6 (Ritchie, Phipson et al. 2015), in R, version 4.4.0^67^. Gene set enrichment analysis (GSEA) was performed using WebGestaltR version 0.4.6^68^. Clustering and visualization of the GSEA results was done using aPEAR version 1.0.0^69^.

### Gene Ontology

All proteomics data sets were analyzed by ShinyGO 0.80.

### Total RNA isolation and quantitative PCR

RNA was isolated using the Quick RNA Miniprep kit (Zymo Research, R1054) according to the manufacturer’s protocols. 500 ng total RNA was used as input for the synthesis of cDNA using the ABScript III RT Master Mix (ABclonal, RK20429). qPCRs were performed using the Universal SYBR Green Fast qPCR Mix (ABclonal, RK21203) on a CFX384 Touch Real-Time PCR Detection System (BioRad). Gene expression values were normalized using mouse *Rpl37a1* as a relative fold change to the value of control samples. All experiments were assayed by the ΔΔCt method. All used primers for qPCR analysis are listed in **Supplementary Table 4**.

### Statistical analysis

Data are presented as mean ± SEM/SD. The mean of technical replicates is plotted for each biological replicate. Biological replicates represent different passages of the cells that were seeded on independent days. Statistical significance was calculated using GraphPad Prism (GraphPad Software, San Diego, California). Significance levels are * *p* < 0.05, ** *p* < 0.01, *** *p* < 0.001, **** *p* < 0.0001 versus the respective control.

## Acknowledgements

We thank Drs. Sheng Yi Wu, Khyati Gohil, Yufeng Zhao, Takuya Terai and Yusuke Nasu for their valuable advice. We also thank Dr. Maya Shmulevitz for the preparation of cancer cell lines. This work was supported by grants from Canadian Institutes of Health Research (CIHR; FS154310), Japan Science and Technology Agency (JST) CREST (JPMJCR25T3), Mitsubishi foundation, Japan Society for the Promotion of Science (JSPS; 24H00489) and the Max Planck Society. S.H. was supported by a Research Fellowship for Young Scientists from JSPS (24KJ0642). K.T-Y. was supported by the Uehara memorial postdoctoral fellowship. N.T was supported by Overseas Research Fellowships from the JSPS. We also thank University of Alberta Molecular Biology Services Units (MBSU) for DNA sequencing support, P. Giavalisco, and Y. Hinze from the MPI-AGE metabolomics core facility for metabolomics analysis, L. Schumacher, M. Germer and C. Kukat from the MPI-AGE Flow and imaging core facility for supporting the flow cytometry, cell sorting and ImageStream analysis and I. Atanassov, T. Colby, X. Li and I. Matic from the MPI-AGE proteomics core facility for proteomics analysis.

## Author contributions

K.T-Y., S.H., N.T. and Y.S. designed the study and analyzed the data. S.H., N.T., K.T-Y., B.S. A-M and S.M.A. performed the experiments. M.S.M., I.H. and R.E.C. supervised the experiments. S.H., N.T., R.E.C., K.T-Y. and I.H. wrote the manuscript. All authors discussed the results and commented on the manuscript.

