## Supplementary Information for "Ratiometric Fluorescent Protein Biosensors Reveal Citrate Dynamics and Cellular Heterogeneity"

*Corresponding authors.

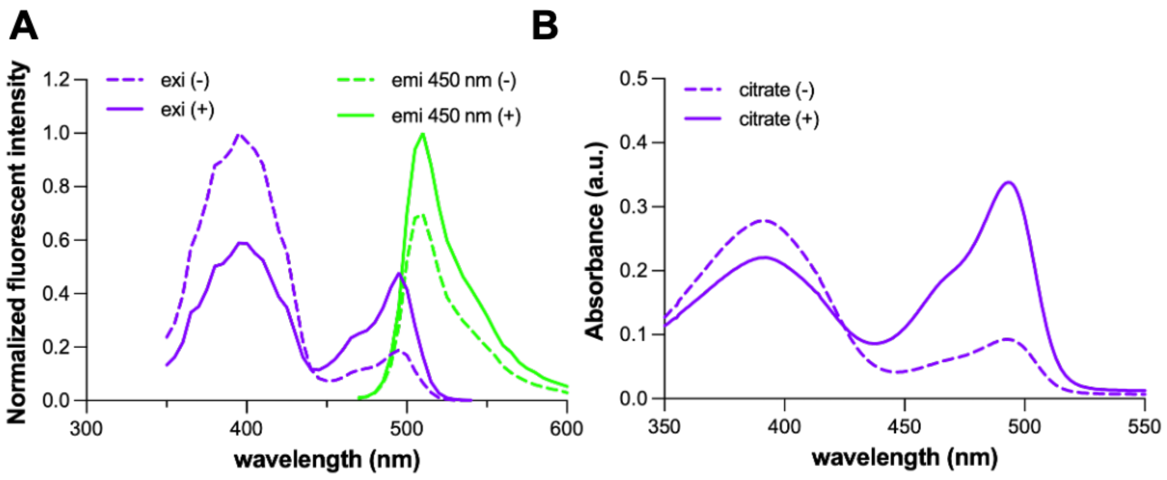

**Supplementary Figure 1. *In vitro* characterization of the initial Citron1-T65A variant.** **A.** Excitation (purple lines) and emission spectra (green lines) of the T65A variant in the absence (dotted lines) and presence (solid lines) of citrate. **B.** Absorbance spectra of the Citron1-T65A variant in the absence (dotted line) and presence (solid line) of citrate.

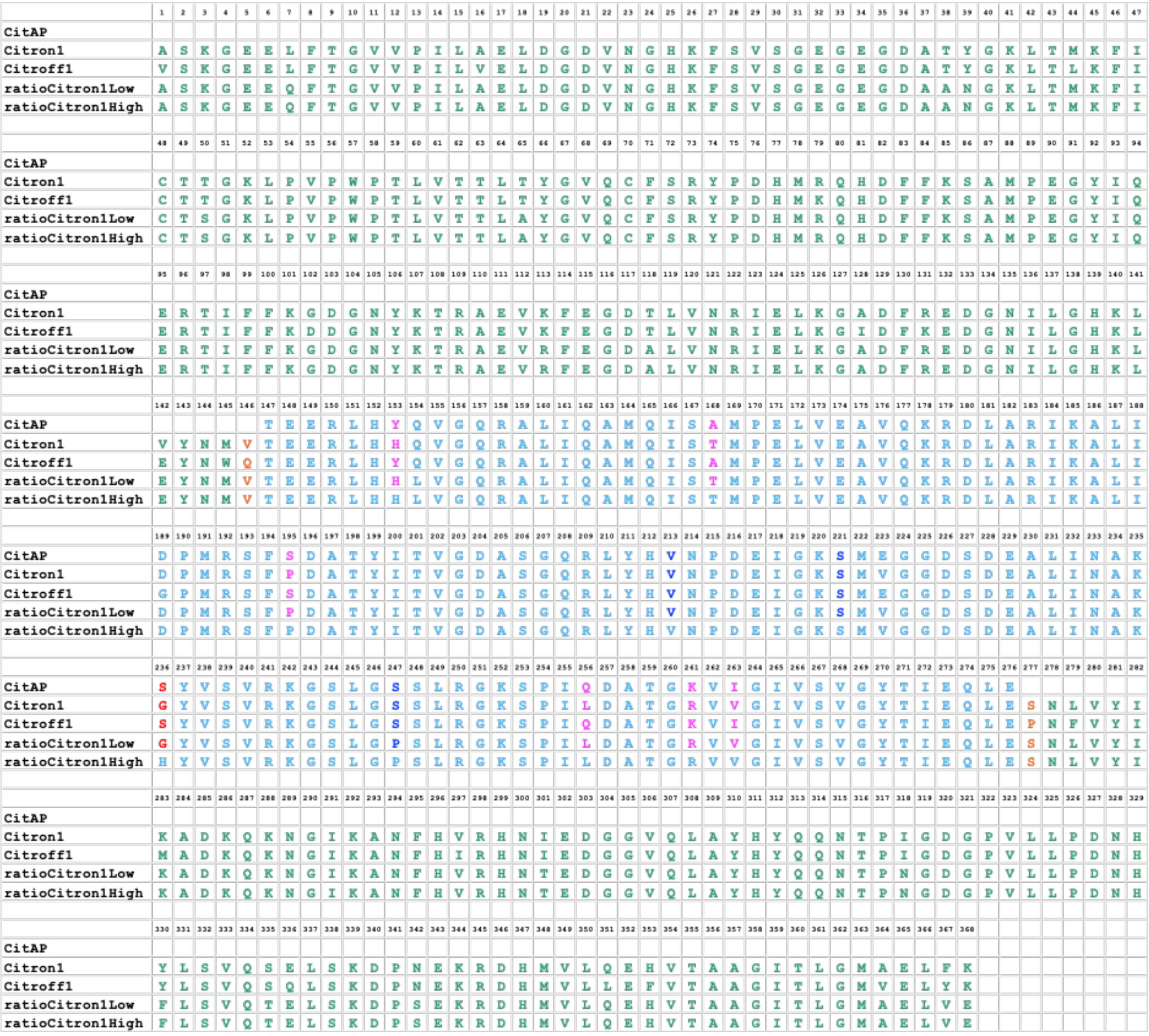

**Supplementary Figure 2. Sequence alignment of CitA and citrate biosensors.**
The sequence alignment of original CitA, Citron1, Citroff1 and ratioCitron1 variants. The GFP sequence is in green, the sensing domain (CitA) sequence is in cyan, and the linker residues are shown in orange. The key residues for citrate binding are in blue. The residues in CitA where mutations are inserted in the development of Citron1 and ratioCitron1 are shown in magenta and red.

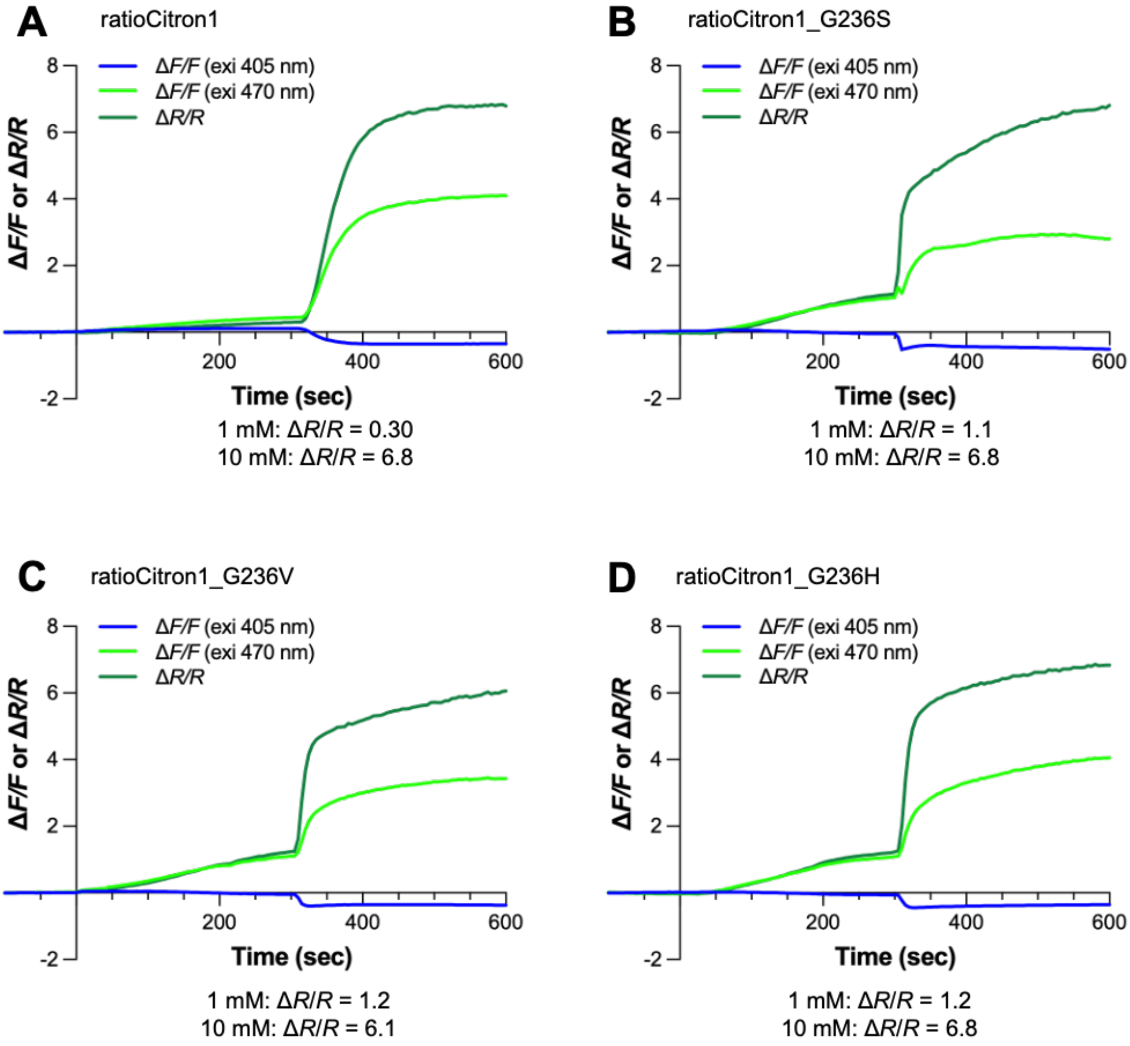

**Supplementary Figure 3. Responses of ratioCitron1 variants to 1 mM and 10 mM citrate in permeabilized HeLa cells.**

**A-D.** Time course of Δ$F/F$ and Δ$R/R$ of ratioCitron1 (**A**), ratioCitron1_G236S (**B**), ratioCitron1_G236V (**C**), and ratioCitron1_G236H (**D**) expressed in HeLa cells. HeLa cells were treated with 4 μM digitonin and a final concentration of 1 mM and 10 mM of citrate was added exogenously at t = 0 and 300 seconds respectively. Δ$F/F$ = ($F_{on}$ - $F_{off}$)/$F_{off}$, and $F$ with excitation 405 nm or 470 nm and emission 518/45 nm, Δ$R/R$ = ($R_{on}$ - $R_{off}$)/$R_{off}$, and $R$ = ($F$ with excitation 470 nm and emission 518/45 nm) / ($F$ with excitation 405 nm and emission 518/45 nm).

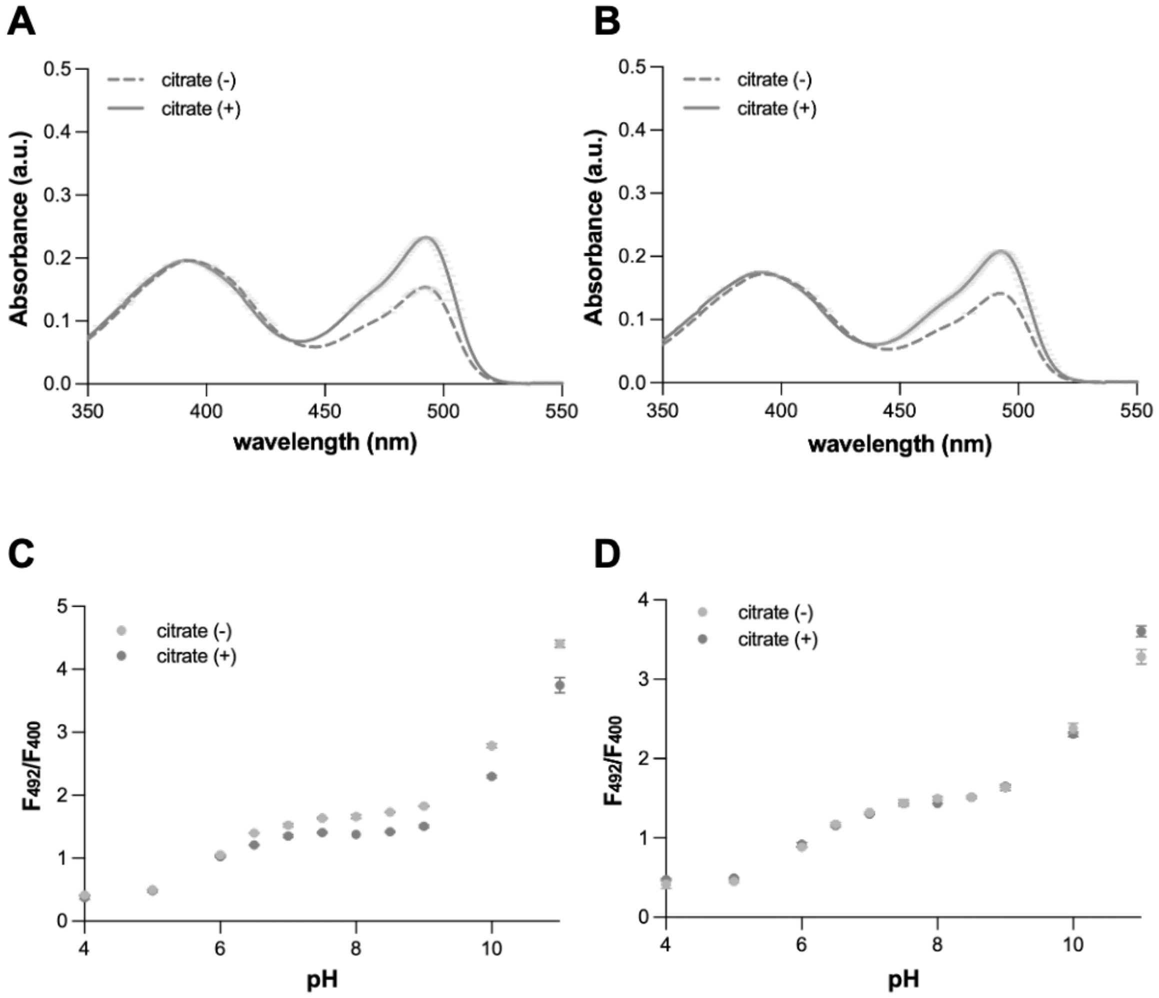

**Supplementary Figure 4. *In vitro* characterization of control variants.**

**A-B.** Absorbance spectra of control variants (ratioCitron1Low-con (**B**) and ratioCitron1High-con (**C**)) in the absence and presence of citrate. *n* = 3 technical replicates, mean ± SD. **C-D.** pH titrations of control variants (ratioCitron1Low-con (**C**) and ratioCitron1High-con (**D**)) in absence and presence of citrate. *n* = 3 technical replicates, mean ± SD.

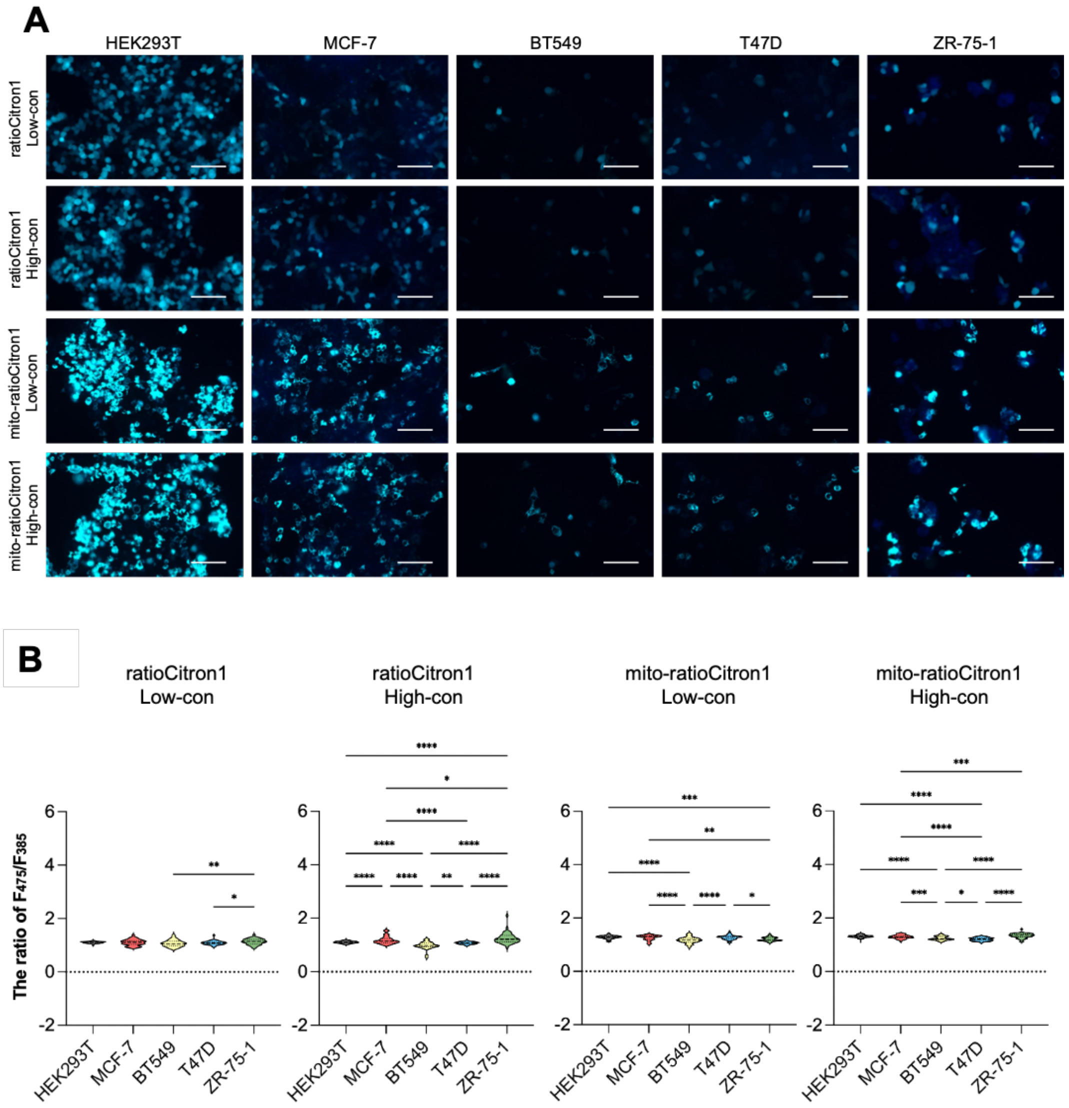

**Supplementary Figure 5. RatioCitron1-control series uncovers no heterogeneity in the ratio of fluorescence from cancer cell lines.**

**A.** Human breast cancer cells and HEK293T cells were transfected with non-responsive ratioCitron1-control variants expressed in cytosol and mitochondria. Representative images are shown. The blue color showed the fluorescent signals taken by 385 nm excitation (emission 499-529 nm) and green color showed the fluorescent signals taken by the 475 nm excitation (emission 500-550 nm). The blue and green overlaid images are shown. Scale bar: 100 μm. **B.** The emission ratio of each excitation wavelength ($F_{475}/F_{385}$) is shown (ratioCitron1Low-con; HEK293T $n$ = 142, MCF-7 $n$ = 53, BT549 $n$ = 39, T47D $n$ = 48, ZR-75-1 $n$ = 47, ratioCitron1High-con; HEK293T $n$ = 121, MCF-7 $n$ = 86, BT549 $n$ = 28, T47D $n$ = 66, ZR-75-1 $n$ = 43, mito-ratioCitron1Low-con; HEK293T $n$ = 110, MCF-7 $n$ = 71, BT549 $n$ = 28, T47D $n$ = 34, ZR-75-1 $n$ = 31, mito-ratioCitron1High-con; HEK293T $n$ = 132, MCF-7 $n$ = 84, BT549 $n$ = 44, T47D $n$ = 41, ZR-75-1 $n$ = 49). This experiment was independently repeated three times and one result from the representative experiment is shown. Statistically analyzed using one-way ANOVA with Tukey's multiple comparisons test (*, $p < 0.05$; **, $p < 0.01$; ***, $p < 0.001$; ****, $p < 0.0001$).

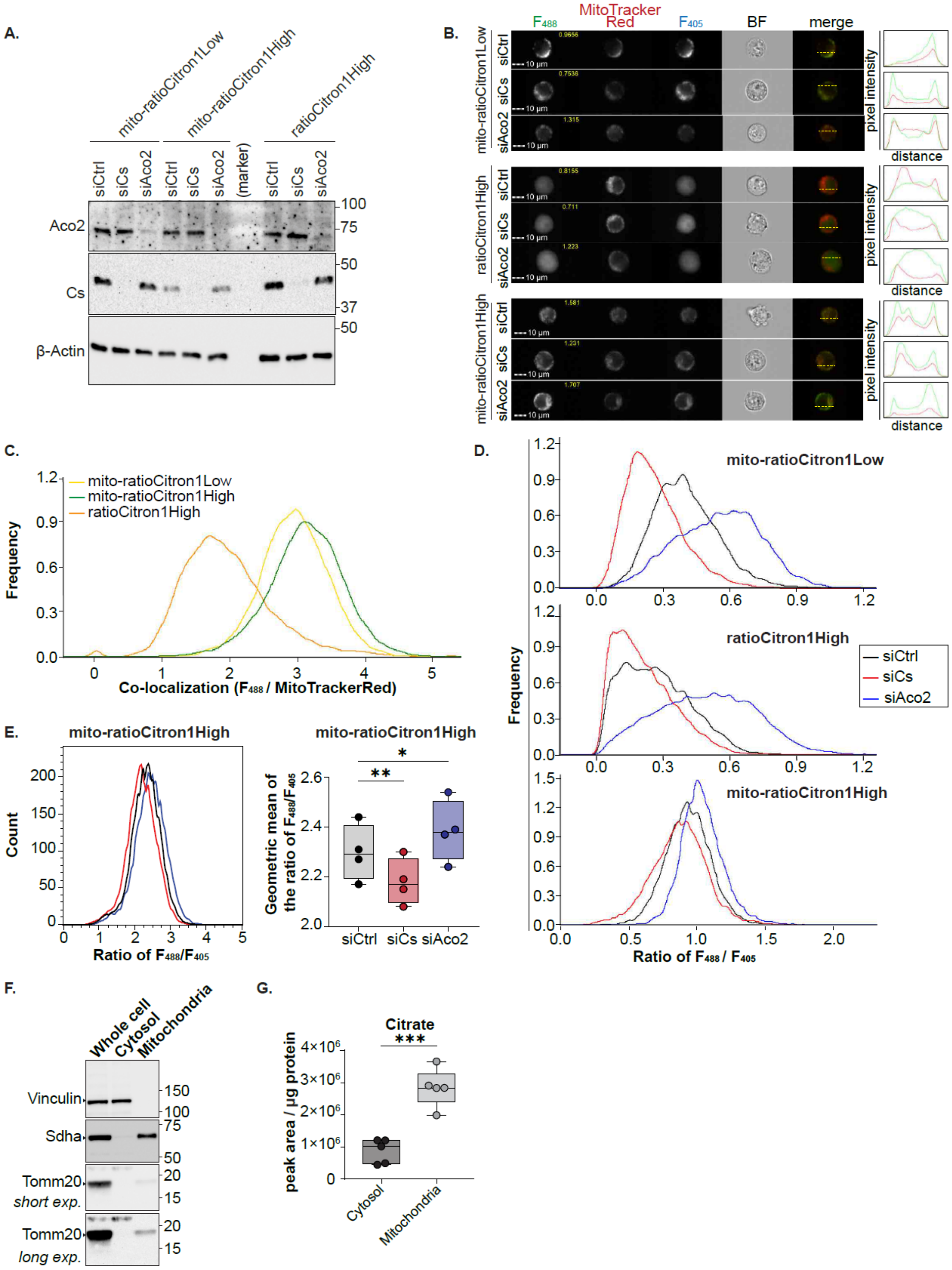

A.
mito-ratioCitron1Low
mito-ratioCitron1High
ratioCitron1High
siCtrl
siCs
siAco2
(marker)
Aco2
Cs
β-Actin
100
75
50
37
50
B.
F488
MitoTracker Red
F405
BF
merge
mito-ratioCitron1Low
ratioCitron1High
mito-ratioCitron1High
pixel intensity
distance
10 µm
C.
mito-ratioCitron1Low
mito-ratioCitron1High
ratioCitron1High
Frequency
Co-localization (F488 / MitoTrackerRed)
D.
mito-ratioCitron1Low
ratioCitron1High
mito-ratioCitron1High
siCtrl
siCs
siAco2
Frequency
Ratio of F488 / F405
E.
mito-ratioCitron1High
Count
Ratio of F488/F405
Geometric mean of the ratio of F488/F405
siCtrl
siCs
siAco2
F.
Whole cell
Cytosol
Mitochondria
Vinculin
Sdha
Tomm20
short exp.
Tomm20
long exp.
150
100
75
50
20
15
G.
Citrate
peak area / µg protein
Cytosol
Mitochondria

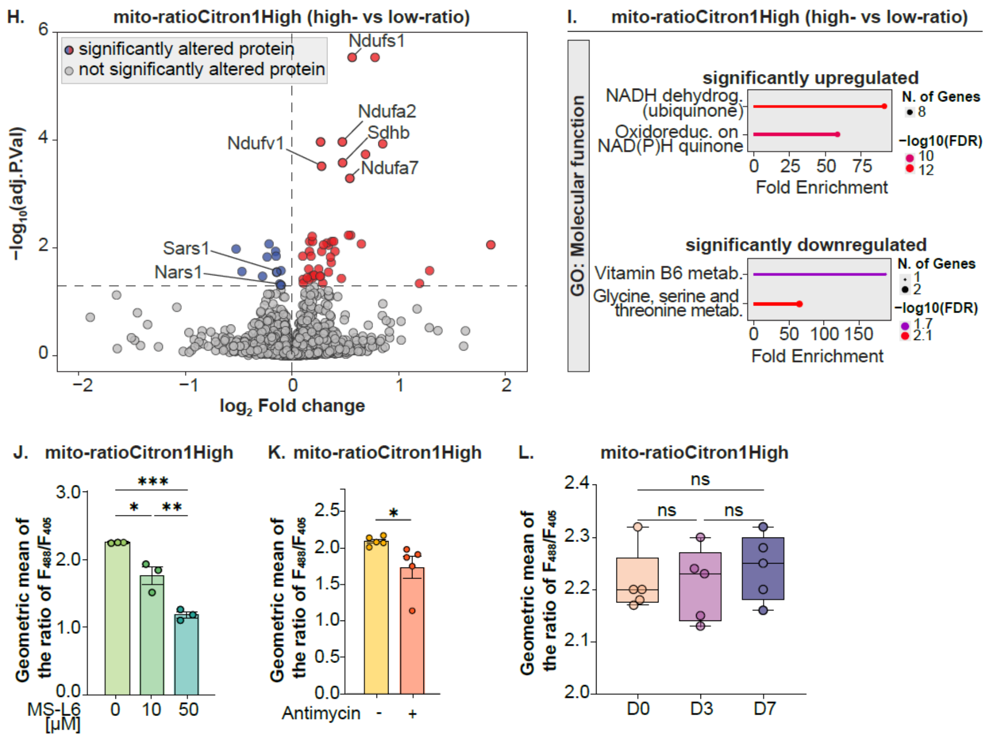

**Supplementary Figure 6. Perturbation of TCA cycle and electron transport chain components modulate mitochondrial citrate heterogeneity.**

**A.** Western blotting analysis of the indicated ratioCitron1 variant KI mESCs after treatment with non-targeting or Cs or Aco2-targeting siRNA using Aco2 and Cs specific antibodies. β-Actin was used as a loading control. **B.** Representative ImageStream analysis of the indicated ratioCitron1 variant KI mESCs upon each siRNA treatment. Fluorescence intensity profiles were obtained from representative images. The $F_{488}/F_{405}$ ratio of representative images is shown in yellow. $n$ = 3 biological replicates. **C.** Frequency of co-localization signals between the $F_{488}$ of the ratioCitron1 variants and MitoTracker Red in non-targeting siRNA-treated mESCs is depicted. **D.** Representative histograms of the $F_{488}/F_{405}$ ratio using cells from (**B**) as input analyzed by ImageStream. Mito-ratioCitron1Low (yellow), mito-ratioCitron1High (green) and ratioCitron1High (orange) data are shown. **E.** mito-ratioCitron1High KI mESCs were treated with siRNA against siCtrl (gray), siCs (red) and siAco2 (blue), respectively and analyzed by flow cytometry. Representative histogram and geometric mean of the $F_{488}/F_{405}$ ratio is shown. $n$ = 4 biological replicates. The box plots represent the median and 25–75 percentiles. Statistically analyzed using one-way ANOVA with Tukey's multiple comparisons test (*, $p$ < 0.05; **, $p$ < 0.01). **F.** Whole cell and fractionated cell lysates were subjected to Western blotting analysis of the cytosolic protein, Vinculin and the mitochondrial proteins, Sdha and Tomm20. **G.** Metabolic measurement of citrate level in cytosolic and mitochondrial fractions of wild-type mESC cells. $n$ = 5 technical replicates. The box plots represent the median and 25–75 percentiles. Statistically analyzed using Student's *t*-test (*** $p$ < 0.001). **H.** Proteomics analysis of the differentially expressed proteins in the high- and low-ratio mito-ratioCitron1High KI cell populations. Significant up- (red) and downregulated (blue) proteins are highlighted. The proteins related to electron transport chain complex (ETC) in the upregulated and aminoacyl-tRNA syntheses in the downregulated are labelled. **I.** Gene ontology analysis of significantly up- or downregulated proteins as shown in (**H**). **J–K.** mito-ratioCitron1High KI mESCs were cultured in the absence or presence of 10 or 50 μM MS-L6 for 1 h (**J**), or 5 nM Antimycin for 2 h (**K**) and analyzed by flow cytometry. Geometric mean of the $F_{488}/F_{405}$ ratio is shown and statistically analyzed using Student's *t*-test or one-way ANOVA with Tukey's multiple comparisons test (*, $p < 0.05$; **, $p < 0.01$; *** $p < 0.001$). $n = 3$ (**J**) and $n = 6$ (**K**) biological replicates. Error bars indicate the SEM. **L.** mito-ratioCitron1High KI mESCs were cultured in the presence or absence of LIF from D0, D3 and D7. Geometric mean of the $F_{488}/F_{405}$ ratio during differentiation is shown and statistically analyzed using or one-way ANOVA with Tukey's multiple comparisons test (ns, not significant). $n = 5$ biological replicates. The box plots represent the median and 25–75 percentiles.

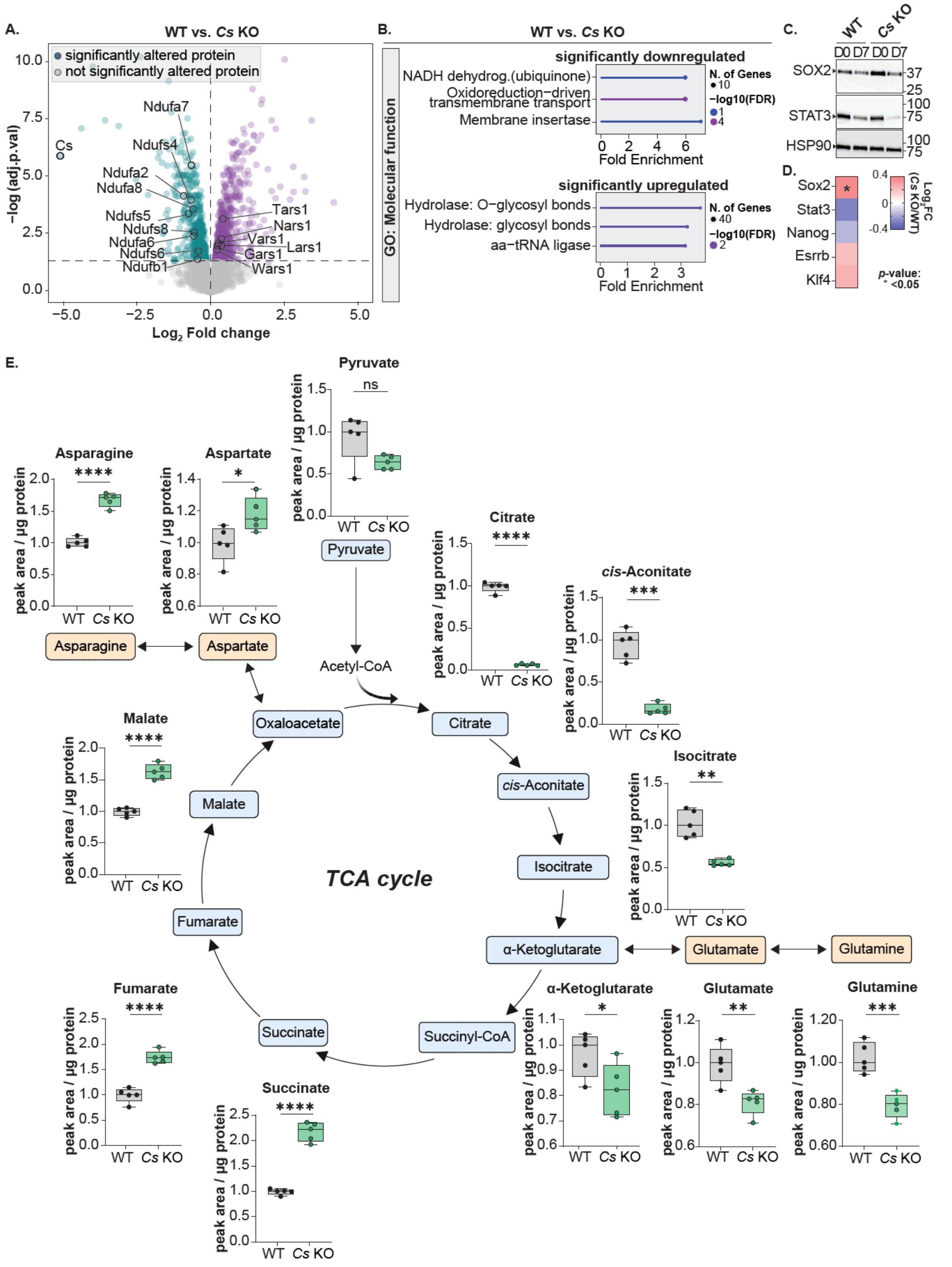

**Supplementary Figure 7. Loss of citrate synthase remodels the proteome and mitochondrial metabolism in pluripotent stem cells.**

**A.** Volcano plot representing the differentially expressed proteins when comparing WT and *Cs* KO mESCs. *n* = 5 biological replicates. Significantly up- (purple) and downregulated (green) proteins are highlighted. Selected proteins related to the electron transport chain (ETC) and Cs in the downregulated fraction and aminoacyl-tRNA synthases in the upregulated fraction are labelled. **B.** Gene ontology analysis of significantly up- or downregulated proteins as shown in (**A**). **C.** Western blotting of D0 and D7 of WT and *Cs* KO mESCs using the pluripotent marker proteins, Stat3 and Sox2, was performed. Hsp90 was used as a loading control. **D.** Comparison of indicated pluripotent markers between WT and *Cs* KO mESCs as in (**A**). Statistically analyzed using Student's *t*-test ($p < 0.05$). **E.** Comparison of indicated steady-state metabolites between WT and *Cs* KO mESCs. $n = 5$ technical replicates. The box plots represent the median and 25–75 percentiles. Statistically analyzed using Student's *t*-test (*, $p < 0.05$; **, $p < 0.01$; ***, $p < 0.001$; ****, $p < 0.0001$).

**Supplementary Table 4: Primer List**

| **Primer for *Cs* KO** | **Sequence (5′ → 3′)** |
|---|---|
| **Single guide RNAs** | |
| *Cs*_sgRNA_fwd | GCAGCAACATGGGAAGACAG |
| *Cs*_sgRNA_rev | CTGTCTTCCCATGTTGCTGCC |
| **Genotyping PCR** | |
| *Cs*_genome PCR_fwd | CATGCCTGGCAGACCAAGAAAC |
| *Cs*_genome PCR_rev | CAGAGGCGAATGAGGATAGGC |
| **qPCR primers** | |
| **taken from Kroef et al**[61] | |
| *Rpl37a*_qPCR_fwd | GTGGTTCCTGCATGAAGACAGTG |
| *Rpl37a*_qPCR_rev | TTCTGATGGCGGACTTTACCG |
| *Fgf8*_qPCR_fwd | AACTCGGACTCTGCTTCCAA |
| *Fgf8*_qPCR_rev | CAGGTCCTGGCCAACAAG |
| *Otx2*_qPCR_fwd | CGTGGGCTACCCCGCCAC |
| *Otx2*_qPCR_rev | CTGGGTACCGGGTCTTGGCA |
| *Sox17*_qPCR_fwd | GAATCCAACCAGCCCACTGA |

| | |
|---|---|
| *Sox17*_qPCR_rev | TAGGGAAGACCCATCTCGGG |
| *Eomes*_qPCR_fwd | GGAAGTGACAGAGGACGGTG |
| *Eomes*_qPCR_rev | AGCCGTGTACATGGAATCGT |
| *Fgf5*_qPCR_fwd | ACCCGGATGGCAAAGTCAA |
| *Fgf5*_qPCR_rev | CAATCCCCTGAGACACAGCAA |
| *Brachyury*_qPCR_fwd | GCTCTAAGGAACCACCGGTCATC |
| *Brachyury*_qPCR_rev | ATGGGACTGCAGCATGGACAG |
| *Hand1*_qPCR_fwd | TCTGGCTCGCTCTCTCGTCC |
| *Hand1*_qPCR_rev | CTCGAGAAGGCATCAGGGTA |
